# Vinculin-Arp2/3 Interaction Inhibits Branched Actin Assembly to Control Cell Migration and Cell Cycle Progression

**DOI:** 10.1101/2023.10.09.561480

**Authors:** John James, Artem I. Fokin, Dmitry Y. Guschin, Hong Wang, Anna Polesskaya, Svetlana N. Rubtsova, Christophe Le Clainche, Pascal Silberzan, Alexis M. Gautreau, Stéphane Romero

## Abstract

Vinculin is a mechanotransducer that reinforces links between cell adhesions and linear arrays of actin filaments upon myosin-mediated contractility. Both adhesions to the substratum and neighboring cells, however, are initiated within membrane protrusions that originate from Arp2/3-nucleated branched actin networks. Vinculin has been reported to interact with the Arp2/3 complex, but the role of this interaction remains poorly understood. Here we compared the phenotypes of vinculin knock-out (KO) cells with those of knock-in (KI) cells, where the point mutation P878A that impairs the Arp2/3 interaction is introduced in the two vinculin alleles of MCF10A mammary epithelial cells. The interaction of vinculin with Arp2/3 inhibits actin polymerization at membrane protrusions and decreases migration persistence of single cells. In cell monolayers, vinculin recruits Arp2/3 and the vinculin-Arp2/3 interaction participates in cell-cell junction plasticity. Through this interaction, vinculin controls the decision to enter a new cell cycle as a function of cell density.

## INTRODUCTION

Actin dynamics control cell shape, adhesion and migration (Pollard & Cooper, 2009). Actin filaments form linear and branched arrays (Le Clainche & Carlier, 2008). Whereas branched actin arrays generate pushing forces through the Arp2/3 complex, linear arrays can generate pulling forces through myosin-mediated contractility (Pollard, 2016; Garrido-Casado *et al*, 2021). Indeed, cell migration requires a combination of forces to drive protrusion of the plasma membrane and to pull on the substratum (Blanchoin *et al*, 2014).

The RAC1-WAVE-Arp2/3 pathway drives membrane protrusions (Papalazarou & Machesky, 2021; Bieling & Rottner, 2023). This pathway is embedded in positive feedback loops that sustain the membrane protrusion at the front edge of a migrating cell and increases the persistence of migration (Krause & Gautreau, 2014). Signaling from branched actin at the cell cortex is also critical for cells to progress into the cell cycle by impinging on tumor suppressor genes controlling the G1/S transition (Molinie *et al*, 2019). Various Arp2/3 complexes coexist in the same cells through the combinatorial assembly of paralogous subunits (Pizarro-Cerdá *et al*, 2016). For example, ARPC1B-containing Arp2/3 complexes are more efficient at nucleating branched actin and at forming stable branched junctions than ARPC1A- containing Arp2/3 complexes (Abella *et al*, 2016). ARPC1B-containing Arp2/3 complexes, but not the ones containing ARPC1A, generate cortical branched actin that drives persistent migration and delivers a signal for cell cycle progression (Molinie *et al*, 2019).

Vinculin is recruited by mechanosensitive proteins that sense forces exerted in cell adhesions and responds to these forces by reinforcing the linear arrays of actin filaments attached to cell adhesions (Bays & DeMali, 2017). Vinculin is composed of a head that interacts with adhesion proteins and a tail that binds to actin filaments (Humphries *et al*, 2007; Le Clainche *et al*, 2010). This ability of vinculin to link actin filaments to adhesion sites is inhibited by an intra-molecular interaction masking relevant binding sites of the head and the tail (Atherton *et al*, 2016). In adhesions to the extracellular matrix (ECM), vinculin head is recruited to cryptic binding sites in talin that are exposed upon stretching due to myosin-mediated contractility (del Rio *et al*, 2009; Ciobanasu *et al*, 2014; Yao *et al*, 2014a). Vinculin head is similarly recruited to a cryptic binding site of α-catenin at cell-cell adhesions upon myosin-mediated contractility of actin filaments (Yao *et al*, 2014b; Seddiki *et al*, 2018; Vigouroux *et al*, 2020). Force-dependent recruitment of vinculin results in its activation allowing vinculin tail to link additional actin filaments and thereby reinforce the cytoskeletal attachment of cell adhesions. In the process, vinculin mechanotransduces signals, since vinculin-depleted cells exhibit enhanced proliferation, survival and anchorage-independent growth (Fernández *et al*, 1993; Subauste *et al*, 2004; DeWane *et al*, 2022).

The cytoskeleton reinforcement function of vinculin is in line with the fact that cell adhesions mature over time. Adhesions to the ECM mature from nascent adhesions at the edge of membrane protrusions into focal adhesions (FAs) as the leading edge moves forward and myosin motors exert contractility on newly formed adhesions (DePasquale & Izzard, 1991; Alexandrova *et al*, 2008; Thievessen *et al*, 2013). Similarly, E-Cadherin based adherens junctions (AJs) form initial interdigitations that mature into straight cell-cell adhesions, as contractility develops (Kovacs *et al*, 2002; Leerberg *et al*, 2014; Li *et al*, 2020, 2021). Vinculin was shown in vitro to remodel branched actin networks into bundles (Boujemaa-Paterski *et al*, 2020). In this context, the reported interaction of vinculin with Arp2/3, the major player in protrusion formation, is particularly intriguing.

Vinculin binds to the Arp2/3 complex through the linker that connects its head to its tail (DeMali *et al*, 2002). This linker is not masked by the head-to-tail intramolecular interaction (Borgon *et al*, 2004), suggesting that the interaction should be independent from vinculin activation. Subsequently, ‘vinculin-Arp2/3 hybrid complexes’ were purified from chicken smooth muscles (Chorev *et al*, 2014). These hybrid complexes lack some subunits of the canonical Arp2/3, namely ARPC1, ARPC4 and ARPC5 subunits. These observations suggest the existence of constitutive vinculin-Arp2/3 complexes. However, the interaction of vinculin with Arp2/3 is regulated by EGF stimulation, RAC1 activity and Src-dependent phosphorylations of vinculin on tyrosine residues that are located far away from the Arp2/3 binding site, but which contribute to vinculin activation (DeMali *et al*, 2002; Zhang *et al*, 2004; Moese *et al*, 2007; Auernheimer *et al*, 2015). The point mutation P878A in vinculin was shown to impair its interaction with Arp2/3 and a re-expression of this mutant in vinculin-null mouse embryonic fibroblasts (MEFs) does not rescue defective protrusions and spreading of these cells, unlike wild type vinculin (DeMali *et al*, 2002). Together these data suggest that vinculin could activate Arp2/3 and form membrane protrusions.

Here we used epithelial cells to investigate the role of the vinculin-Arp2/3 interaction. We were able to distinguish the specific subset of functions that the vinculin-Arp2/3 interaction endows, compared with the more general role of vinculin in the cytoskeletal reinforcement of cell adhesions. We found that the vinculin-Arp2/3 interaction inhibits actin assembly in membrane protrusions, and thereby antagonizes branched actin in the control of cell migration, cell cycle progression, and regulates at cell-cell junction plasticity through Arp2/3 recruitment.

## RESULTS

### The Vinculin Linker Enhances Membrane Protrusion and Migration Persistence through Arp2/3 Binding

We first examined the interaction of vinculin with the Arp2/3 complex in the MCF10A cell line, where we have characterized the role of the Arp2/3 complex in cell migration (Molinie *et al*, 2019; Simanov *et al*, 2021). MCF10A cells are human mammary epithelial cells, which are immortalized but not transformed (Soule *et al*, 1990). Importantly, this cell line is diploid for the most part of its genome (Worsham *et al*, 2005; Kadota *et al*, 2010). We found that antibiotic-selected MCF10A cells down-regulated the exogenous expression of tagged full-length vinculin in an increasing number of cells over time, but not when the construct was limited to the vinculin linker that connects the head to the tail and which contains the Arp2/3 binding site. We obtained stable MCF10A cell lines expressing the GFP tagged linker of vinculin (amino acids 811-881) or its derivative containing the point mutation P878A that impairs Arp2/3 binding (DeMali *et al*, 2002). When mutated, the linker indeed bound much less efficiently to the Arp2/3 complex despite an expression level similar to the wild type linker (Fig. 1a). Interestingly, the immunoprecipitate of the vinculin linker contained the Arp2/3 subunit ARPC1B, which is not present in the vinculin-Arp2/3 hybrid complexes that were purified from tissues (Chorev *et al*, 2014), suggesting that in our cell system, vinculin binds to the canonical Arp2/3 complex.

**Figure 1.**
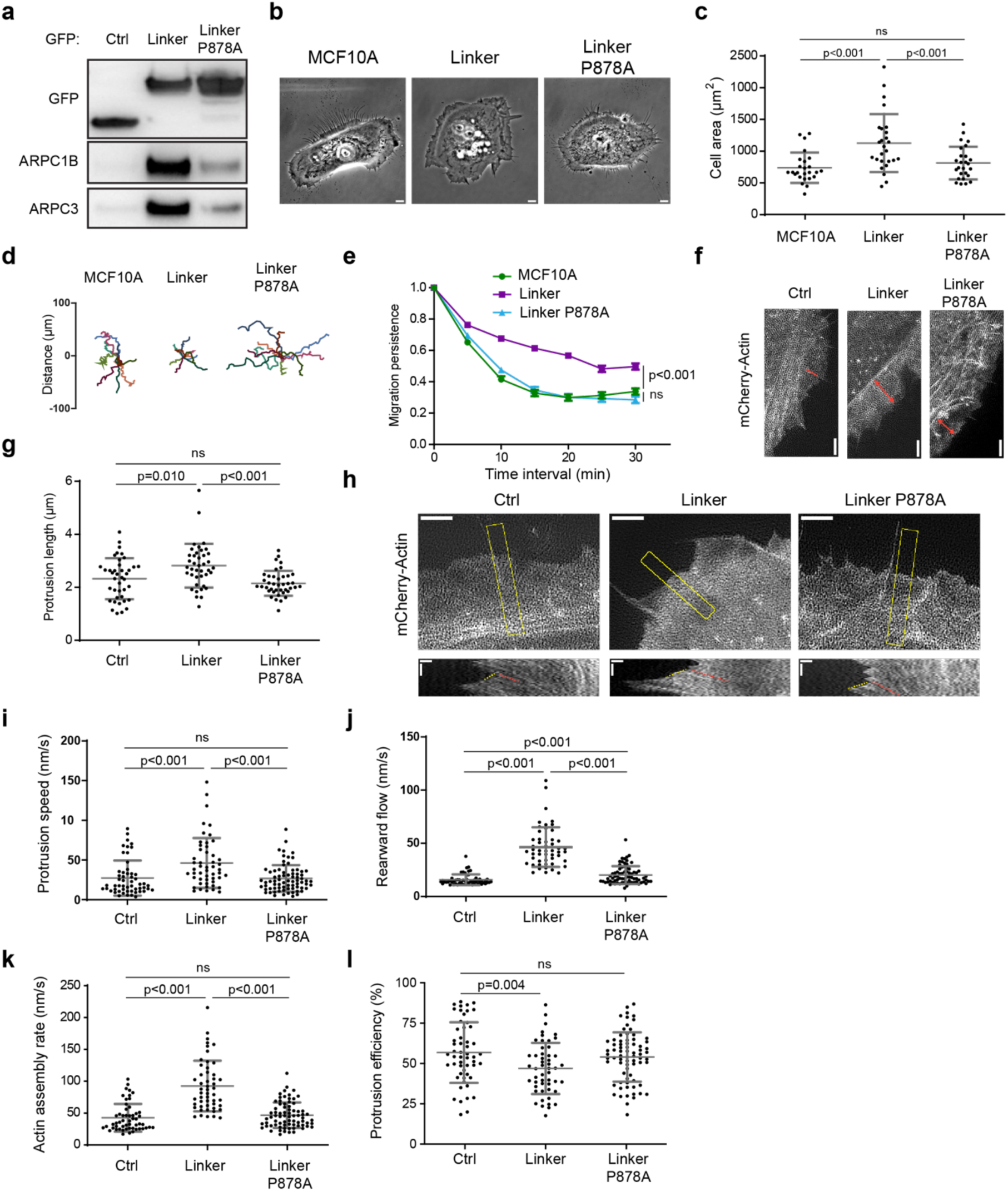
Expression of the vinculin linker that binds to the Arp2/3 complex increases actin polymerization and membrane protrusion. **a** The Arp2/3 complex co-immunoprecipitates with the vinculin linker (amino acids 811-881). MCF10A cells stably expressing GFP, the GFP tagged linker in a WT or P878A form were lysed and subjected to GFP immunoprecipitation and Western blot analysis. N=3, 1 representative experiment shown. **b** Phase contrast images of the same cell lines. Scale bar = 5 μm. **c** Cell area of the cell lines stained for actin. Mean ±SD, n=25, t-test. N=3, 1 representative experiment shown. **d, e** Single cell migration. Cell trajectories **(d)** and migration persistence (**e**). Mean ± SD, n=10, linear mixed effect model. N=3, 1 representative experiment shown. **f, g** Membrane protrusions of the stable MCF10A cell lines expressing the vinculin linker transiently transfected with mCherry-actin. TIRF-SIM images of mCherry-actin **(f)** and quantification of protrusion length **(g)**. Double-headed arrows in red indicate the length of lamellipodia. Scale bar 2 μm. Mean ± SD, n=40, t-test. N=3 with similar results, pooled measurements from the 3 independent repeats are plotted. **h** Kymographs (bottom panels, scale bars 0.4 µm horizontal, 40 s vertical) were generated along a line centered in the region boxed in yellow in the TIRF SIM movie (scale bar 2 μm). Dashed yellow and red lines indicate protrusion speed **(i)** and rearward flow **(j)**, respectively. Actin assembly rate **(k)** is the sum of protrusion speed and rearward flow. Protrusion efficiency **(l)** is the ratio of protrusion speed to actin assembly rate. Mean± SD, n= 40, t-test. N=3 with similar results, pooled measurements from the 3 independent repeats are plotted.

MCF10A cells expressing the wild type linker, but not its P878A derivative, exhibited extensive membrane protrusions at their periphery (Fig. 1b, Supplementary Movie 1). Cells expressing the vinculin linker were significantly more spread (Fig. 1c) and migrated more persistently than parental cells (Fig. 1d,e), two read-outs of cortical polymerization of branched actin. Decreased cell speed and mean square displacement (MSD) were associated with this increased persistence of cell migration (Supplementary Fig.1). Decreased speed and MSD are often associated with increased cortical branched actin in MCF10A cells, for example when RAC1 is activated by mutation or when the Arp2/3 inhibitory protein Arpin is down-regulated (Molinie *et al*, 2019). In contrast, cell migration persistence systematically correlates with increased polymerization of cortical branched actin (Dang *et al*, 2013; Molinie *et al*, 2019; Simanov *et al*, 2021).

We then examined membrane protrusions that power cell migration. The extension of membrane protrusions depends on the efficiency of the molecular clutch that couples cell adhesions to branched actin polymerizing against the plasma membrane (Thievessen *et al*, 2013; Hirata *et al*, 2014). Thus, the speed of membrane protrusions is a function of both rate of actin assembly and actin rearward flow. Following mCherry-actin expression, we were able to measure speed as well as the length of the protrusions, defined as the distance between the protrusion edge and the first transverse arc at the base of protrusions (Fig. 1f). Protrusions were longer and protruded 1.7-fold faster when cells expressed the wild type vinculin linker, but not the P878A derivative (Fig. 1g,h,i). Fluorescent actin also allowed us to image the rearward flow of the cytoskeleton using Total Internal Reflection Fluorescence – Structured Illumination Microscopy (TIRF-SIM, Supplementary Movie 2). We found that both the rearward flow, relative to the substratum, and the actin assembly rate, relative to the leading edge, were significantly increased in cells expressing the vinculin linker (Fig. 1h,j,k). However, protrusion efficiency, the ratio of protrusion speed to actin assembly rate, was slightly decreased in cells expressing the vinculin linker, indicating that protrusion speed does not fully scale with actin assembly rate (Fig. 1l). This effect on protrusion efficiency depended on the interaction between the vinculin linker and Arp2/3 since the expression of the P878A derivative does not modify actin assembly rate, nor protrusion speed, compared with control cells (Fig. 1h,j,k,l). Together, these results suggest either that vinculin activates the Arp2/3 complex through the linker domain, or that vinculin inhibits Arp2/3 and the linker behaves as a dominant-negative fragment. Therefore, we produced and purified the vinculin linker in its wild type and P878A forms. In vitro, these proteins did not modify the kinetics of spontaneous actin polymerization, nor Arp2/3-mediated branched nucleation (Supplementary Figure 2), indicating that the observed effects of the vinculin linker in cells described above require additional factors or post-translational modifications that are only present in cells in order to regulate Arp2/3.

### Vinculin Knock-In of the Point Mutation that Impairs Arp2/3 Interaction Increases Actin Dynamics and Migration Persistence as Vinculin Knock-Out Does

Since no activity of the linker can be revealed in vitro, we generated knock-outs of the *VCL* gene that encodes vinculin in order to distinguish between the linker-mediated activation and dominant-negative hypotheses. We transfected MCF10A cells with purified Cas9 and a guide RNA (gRNA) that targets the beginning of the Open Reading Frame (ORF) in the first exon of the *VCL* gene. Cas9-mediated double strand breaks (DSBs) are frequently repaired by the error-prone non-homologous end joining (NHEJ) mechanism (Jasin & Haber, 2016). This method using purified Cas9 and gRNA was sufficiently efficient to identify KO clones by Western blot-based screening without selection (Fig. 2a). We identified two KO clones with different frame shifts within the two alleles of *VCL* (Fig. 2b), for further characterization. As expected, focal adhesions (FAs) of these two clones were not stained by vinculin antibodies (Fig. 2c,d). Paxillin-stained FAs were significantly more elongated, by 60 % on average, in vinculin KO compared with parental cells (Fig. 2e).

**Figure 2.**
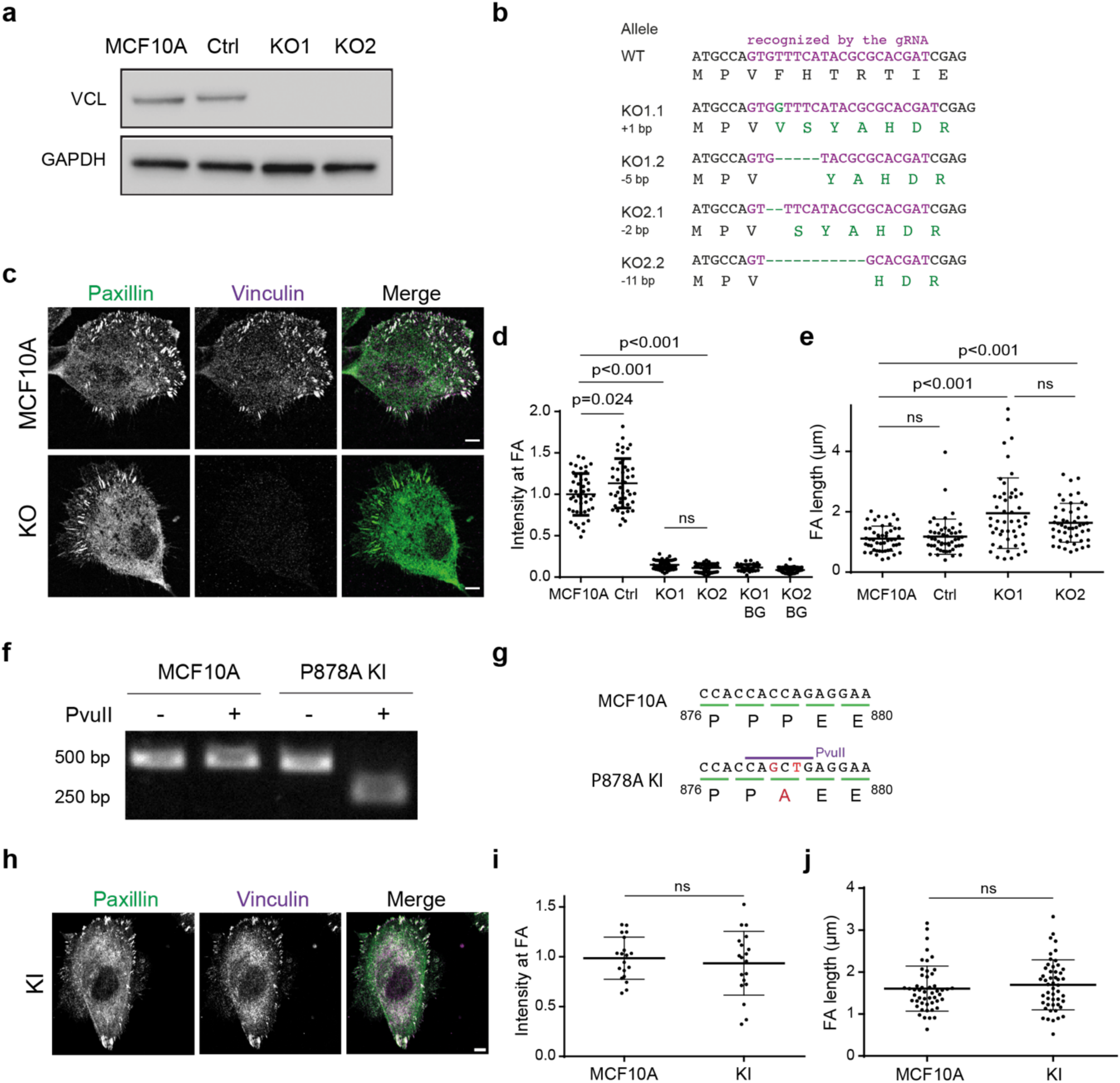
Characterization of *VCL* knock-out and knock-in cell lines. **a** Parental MCF10A, and isolated clones transfected with *VCL* targeting gRNA or a non-targeting gRNA were analyzed by Western blot. **b** Sequences of the two alleles in each KO cell line. All mutations induce a frameshift and thus a premature stop codon in the ORF. **c** Staining of vinculin and paxillin in parental and KO cells. Scale bar = 5 μm. **d** Quantification of vinculin staining in focal adhesions (FAs) and normalization by the intensity of parental cells. BG refers to the background in the non-FA cytoplasm. Mean ± SD, n=45, t-test. N=3 with similar results, pooled measurements from the 3 independent repeats are plotted. **e** Quantification of length of FAs. Mean ± SD, n=45, t-test. N=3 with similar results, pooled measurements from the 3 independent repeats are plotted. **f** Genome analysis of the KI. Part of the *VCL* ORF containing the P878A mutation was amplified by PCR and digested with PvuII restriction enzyme. Agarose gel electrophoresis of digested or undigested PCR fragment. **g** Sequencing of the genome amplified PCR fragment confirmed the presence of the P878A mutation on the two alleles and the introduction of the PvuII restriction site in the genome of the KI line. **h** Staining of vinculin and paxillin in the KI cells. Scale bar = 5 μm. **i** Quantification of vinculin staining in focal adhesions (FAs) and normalization by the intensity of parental cells. Mean ± SD, n=19, t-test N=3 with similar results, pooled measurements from the 3 independent repeats are plotted. **j** Quantification of length of FAs. Mean ± SD, n=50, t-test N=3 with similar results, pooled measurements from the 3 independent repeats are plotted.

To specifically examine the role of the Arp2/3 interaction, we designed a strategy based on homology-directed repair (HDR) to introduce the P878A point mutation in the endogenous *VCL* gene of MCF10A cells. We transfected MCF10A cells with 3 plasmids and a long repair oligonucleotide that provides the P878A mutation together with a PvuII restriction site. One of the plasmids provided Cas9, the second plasmid encoded a gRNA that allows Cas9 to cut the *VCL* gene close to the P878 codon and the third one encoded a gRNA that targets the *ATP1A1* gene. *ATP1A1* encodes the ubiquitous and essential sodium-potassium ion pump, which is the target of the ouabain drug. The gRNA targets the ATP1A1 region that is directly recognized by ouabain. Upon ouabain treatment, only cells that have repaired the *ATP1A1* DSB by NHEJ so as to introduce an in-frame indel, produce an ion pump that is both functional and insensitive to ouabain (Agudelo *et al*, 2017). The ouabain selection is more efficient than more classical antibiotic selection of transfected plasmids, because it ensures that Cas9-mediated DSBs are efficiently produced in the selected cells, and not only that the selected plasmid is present. The DSBs in *VCL* are often repaired by NHEJ, but can also be repaired by HDR using the provided oligonucleotide as a template. After extensive screening of clones by PvuII restriction of the PCR amplified genomic region, we managed to isolate the desired knock-in (KI) clone, where the P878A mutation was introduced in both alleles of *VCL* (Fig. 2f,g). P878A vinculin properly localized to FAs (Fig. 2h,i) and did not impact elongation of FAs (Fig. 2j), suggesting that it is functional and that Arp2/3 binding is not required for the FA-related functions of vinculin.

We then evaluated KO and KI clones for their ability to migrate and produce membrane protrusions. Like the MCF10A clone that expressed the vinculin linker, KO clones exhibited more persistent trajectories than parental cells (Fig. 3a,b, Supplementary Movie 3). They were also 20 % more spread than parental cells (Fig. 3e). The KI clone that expresses the vinculin protein containing the P878A substitution displayed a similar, but even more pronounced phenotype of increased persistence and spreading than the KO clones that expressed no vinculin (Fig. 3c-f). This suggests that the vinculin-Arp2/3 interaction is critical for these functions.

**Figure 3.**
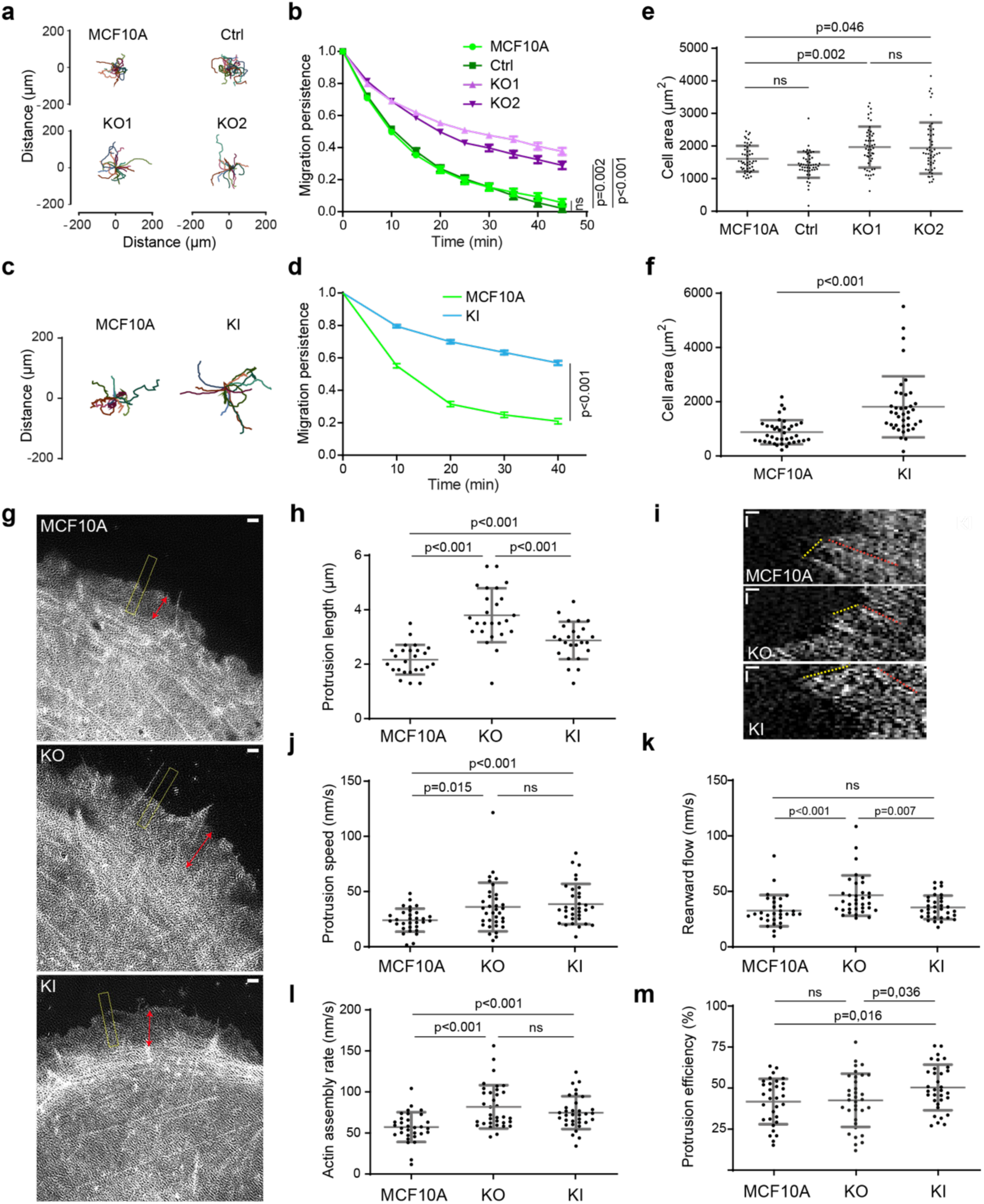
Vinculin decreases migration persistence and actin dynamics in membrane protrusions through its interaction with the Arp2/3 complex. **a-d** Single cell migration of KO and KI cells. Cell trajectories **(a, c)** and migration persistence (**b, d**). Mean ± SD, n=74 in **b** and n=35 in **d** linear mixed effect model. N=3, 1 representative experiment shown. **e, f** Cell area of KO and KI lines. Mean ± SD, n=50 in **e** and n=39 in **f**, t-test. N=3, 1 representative experiment shown. **g, h** Membrane protrusions of the parental MCF10A, KO and KI cell lines transiently transfected with mCherry-actin. TIRF-SIM images of mCherry-actin **(g)** and quantification of protrusion length **(h)**. Double-headed arrows in red indicate the length of lamellipodia and the dashed yellow box indicates the position of kymograph analysis. Double-headed red arrows measure lamellipodia length **(h)**. Scale bar 1 μm. Mean ± SD, n=25, t-test. N=3 with similar results, pooled measurements from the 3 independent repeats are plotted. **i-m** Kymograph analysis of KO and KI lines drawn along a line centered in the region boxed in yellow depicted in panel **g**. Scale bars 0.25 µm horizontal, 10 s vertical **(i)**. Protrusion speed (**j**) is measured from yellow dashed lines and rearward flow **(k)** from red dashed lines. Actin assembly rate **(l)** is the sum of protrusion speed and rearward flow. Protrusion efficiency **(m)** is the ratio of protrusion speed to actin assembly rate. Mean ± SD, n=29, t-test. N=3 with similar results, pooled measurements from the 3 independent repeats are plotted.

We then examined actin dynamics in membrane protrusions by transiently transfecting our KO and KI clones with mCherry-actin. The length of membrane protrusions was increased in both KO and KI clones compared with parental cells (Fig. 3g,h). The increase was, however, significantly higher in KO than in KI cells for this parameter. In KO cells, protrusions extended 1.5-fold faster than in parental cells, in line with a significant increase of actin assembly rate (Fig. 3i,j, l, Supplementary Movie 4). The actin rearward flow was increased in KO cells, indicative of a less efficient molecular clutch in the absence of vinculin (Fig. 3i,k). As a consequence, protrusion efficiency was not different in KO and parental cells (Fig. 3m). In KI cells, protrusions extended and actin assembled as fast as in KO cells (Fig. 3i,j,l, Supplementary Movie 4). But because the clutch was not deficient in this case, the actin rearward flow measured in KI cells was similar to that of parental cells (Fig. 3k), resulting in protrusions of KI cells 1.2-fold more efficient than protrusions of KO and parental cells (Fig. 3m). Together, these results show that the vinculin-Arp2/3 interaction inhibits actin assembly at the leading edge of the lamellipodium, whereas the molecular clutch, which couples branched actin to cell adhesions depends on the presence of vinculin, but not its interaction with Arp2/3. Since the phenotype induced by expression of the vinculin linker is most similar to the vinculin KO phenotype, we can conclude that the linker is dominant negative over the functions of vinculin in the cell.

### Vinculin Controls Stability of Cell-Cell Junctions and Efficiency of Collective Migration

Since vinculin plays a key role in stabilizing cell-cell junctions (Bays & DeMali, 2017), we used our cell system to examine the potential role of vinculin-Arp2/3 interaction in junction stability. We verified that KO cells displayed no vinculin staining at cell-cell junctions (Fig. 4a,b). P878A vinculin expressed in KI cells was recruited at α-catenin positive junctions as efficiently as the wild type protein in parental cells (Fig. 4c,d). Because on rigid 2D substrates such as Petri dishes or glass coverslips, cell-cell interactions were not obviously affected in KO cells, we decided to embed cells into collagen gels to examine cell-cell interactions in a more physiological setting. In 3D collagen gels, behaviors of KO cells and parental cells were markedly different. Parental cells rarely dissociated when they met and formed multicellular slugs (Fig. 4e, Supplementary Movie 5). In contrast, interactions between KO cells appeared not to engage them into a multicellular behavior (Fig. 4e). KI cells did not present the asocial behavior of KO cells and formed multicellular “slugs” like parental cells (Fig. 4f, Supplementary Movie 6). From movies, we were able to count the number of events, where a cell disengaged from cell-cell interactions after having met another cell or dissociating from slugs, per unit of time. The counts were then converted into frequencies expressed in s^-1^ or Hertz. In this 3D setting, the frequency of junction disassembly was about 10-fold higher in KO than parental cells (Fig. 4g). We verified that E-Cadherin expression was not down-regulated in KO cells and that E-Cadherin was properly recruited at cell-cell junctions in KO cells (Supplementary Fig.3). Vinculin thus regulates the stability of junctions, but not E-Cadherin-dependent cell-cell adhesion. In contrast, in KI cells, the quantification revealed that cell-cell junctions were significantly more stable than those of parental cells, dissociating 2.6-fold less (Fig. 4g). Thus, the mechanotransducer function of vinculin is indeed critical to establish stable cell-cell junctions. In contrast, the interaction of vinculin with Arp2/3 is not required for the stability of cell-cell junctions, but rather enhances cell-cell adhesions.

**Figure 4.**
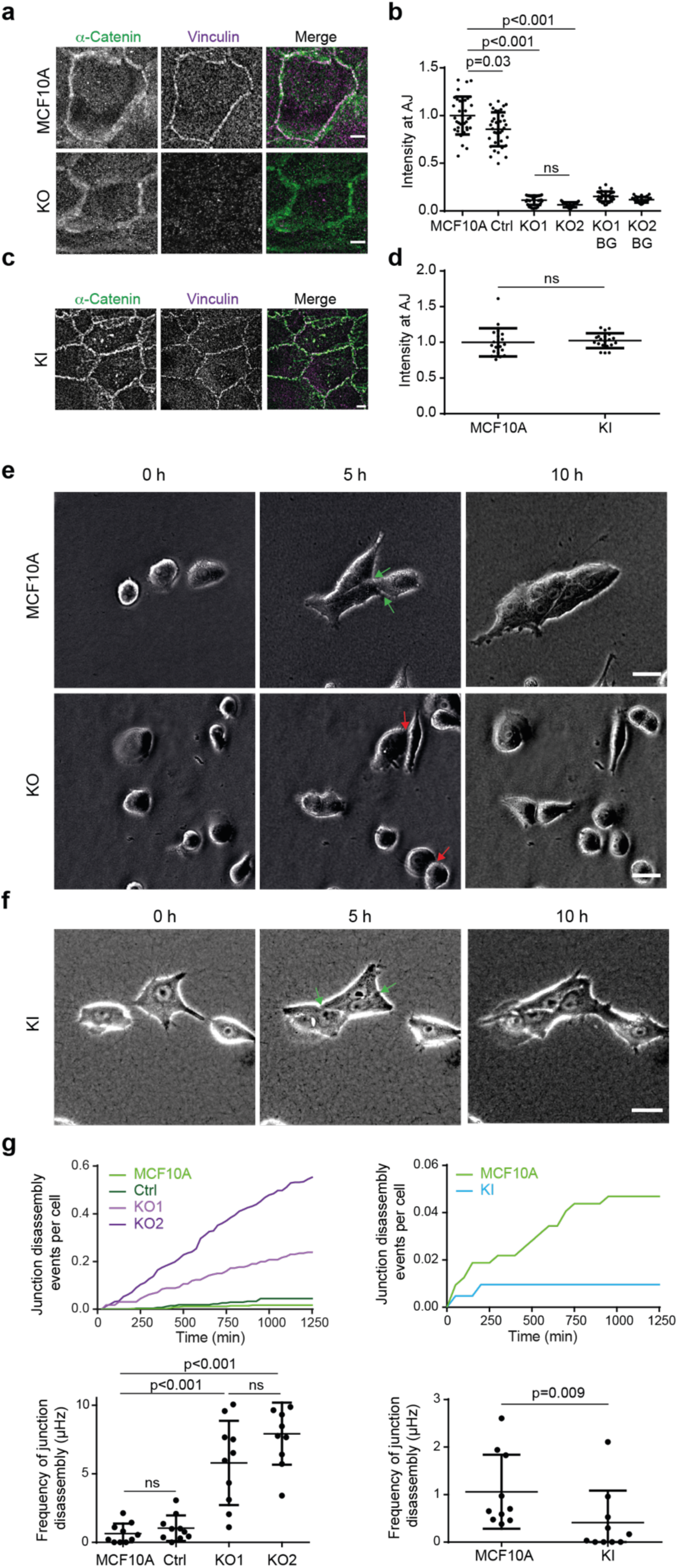
Vinculin stabilizes cell-cell junctions. **a-d** Presence of vinculin at cell-cell junctions in KO and KI cells. Staining of vinculin and α-catenin (**a, c**) and quantification of intensity at cell-cell junctions (**b,d**). Max z-projection from confocal microscopy. Scale bar 5 μm. Mean ± SD, n=37 for **b, n=** 20 for **d**, t-test. N=3 with similar results, pooled measurements from 3 independent repeats are plotted. **e,f** Time-lapse imaging of KO (**e**) and KI (**f**) cells in 3D collagen gels by phase contrast. Green arrows point at cell-cell junctions that were present at 5 h and that did not disassemble at 10 h, red arrows point at cell-cell junctions that were present at 5 h and that were disassembled at 10 h. Scale bars 50 µm in a, 25 µm in b. **g** Quantification of cell-cell junction disassembly events per cell and frequency of cell-cell junction disassembly. Mean ± SD, n=10, t-test. N=3 with similar results, pooled measurements from 3 independent repeats are plotted.

We then analyzed collective migration of MCF10A, KO and KI cells in a wound healing assay obtained by lifting an insert that initially constrains the monolayer (Poujade *et al*, 2007). The front edge of the monolayer of KO cells progressed significantly faster than parental cells and therefore covered the cell-free area more rapidly (Fig. 5a, b, Supplementary Movie 7). KI cells, although slower than KO cells, were also significantly faster than parental cells. Images were then analyzed by Particle Image Velocimetry (PIV) to associate a displacement vector to each (x,y,t) coordinates (Petitjean *et al*, 2010). Cell speed was not only increased at the leading edge of the monolayer in KO and KI cells, but also further backwards, away from the edge, when compared with parental cells (Fig. 5c). The local order parameter, which reflects the local alignment of displacement vectors, was also transmitted further backwards in KO and KI cells when compared with parental cells (Fig. 5d). The increase of both cell speed and local order parameter was further transmitted backwards in KO cells, compared with KI cells, in line with the faster progression of KO front edge. Thus, unlike single cell migration, collective migration partly depends on the mechanotransducer function of vinculin, and partly on its Arp2/3 interaction.

**Figure 5.**
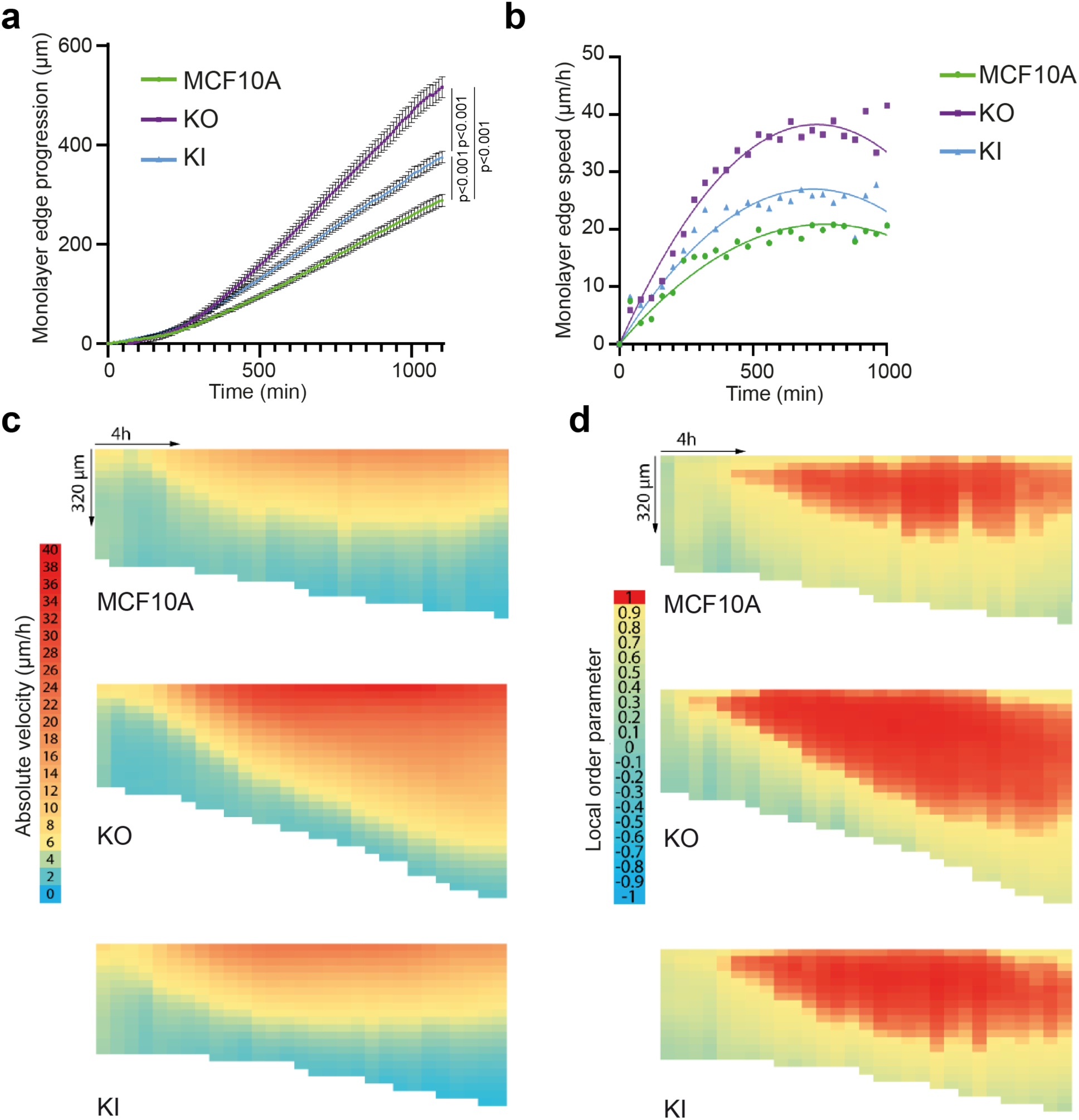
Vinculin controls collective migration upon wound healing. Collective migration of MCF10A, KO and KI cells over the wound was imaged by phase contrast over time and analyzed by Particle Image Velocimetry. **a** Quantification of monolayer edge progression over time. **b** Quantification of monolayer edge speed over time. **c,d** Heat maps representation of velocity (**c**, length of displacement vectors) and local order parameter (**d**, cosine of angles between adjacent displacement vectors). Vertical axis corresponds to coordinates along the perpendicular axis relative to the monolayer edge (edge position kept constant on the top of heat maps), while horizontal axis corresponds to time course (from left to right). N=3 with similar results, 1 representative experiment shown.

### Vinculin Recruits Arp2/3 at Adherens Junctions and Restricts Cell Cycle Progression through Arp2/3 Binding

We then analyzed Arp2/3 recruitment by immunofluorescence at E-Cadherin positive AJs of KO and KI cells. In parental MCF10A cells, Arp2/3 was enriched at AJs, 6 h after plating; the amount of junctional Arp2/3 reached a maximum 1 day after plating and declined to residual amount after 3 days (Fig. 6a). Vinculin was present at AJs throughout junction maturation, even if it also peaked 1 day after plating like Arp2/3 (Supplementary Fig.4). KO cells displayed reduced Arp2/3 staining at AJs compared with parental cells (Fig. 6b). This was especially striking one day after cell plating, when no junctional Arp2/3 was detected. 6 h after plating, a low amount of Arp2/3 was detected at AJs of KO cells, but KO cells still recruited significantly less Arp2/3 than parental cells (Fig. 6b). In KI cells, Arp2/3 recruitment at AJs was increased 6 h after plating compared with parental cells, and even more so compared with KO cells (Fig. 6c,b). Yet, 1 day after plating, Arp2/3 recruitment at AJs of KI cells was dramatically reduced compared with parental cells, as in KO cells. 3 days after plating, Arp2/3 at AJs is minimum in all 3 types of cells.

**Figure 6.**
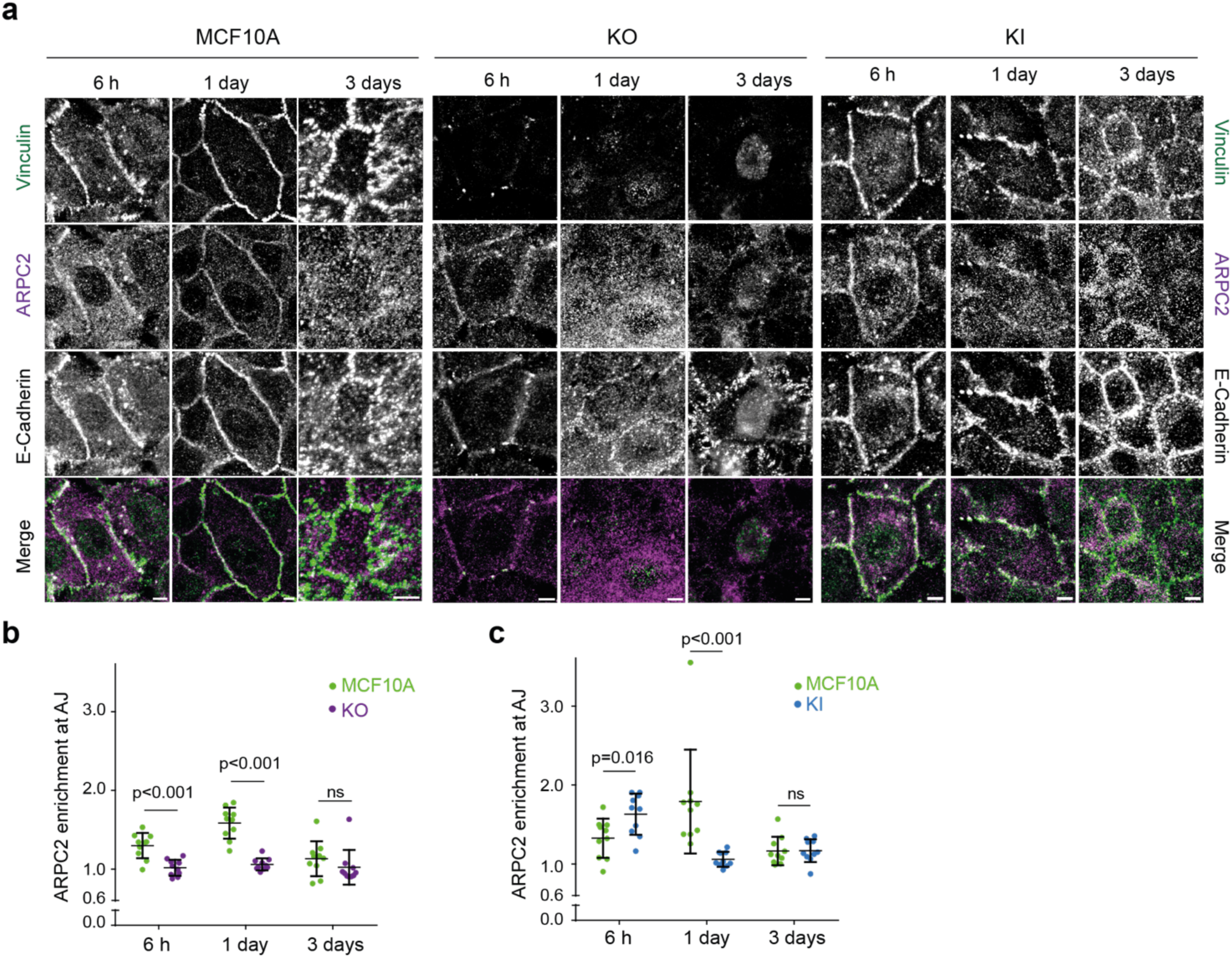
Vinculin retains Arp2/3 at cell-cell junctions. **a** Staining of vinculin, ARPC2 and E-cadherin 6 h, 1 day or 3 days after plating. Scale bar 5 µm. Max z-projection from confocal microscopy. **b,c** Quantification of ARPC2 enrichment at adherens junctions in KO (**b**) and KI (**c**) cells. Mean ± SD, n=10, t-test. N=3, 1 representative experiment shown.

This kinetic analysis thus reveals two distinct phases of Arp2/3 recruitment at AJs. The early recruitment, at 6 h, only partially depends on the presence of vinculin and on its ability to interact with the Arp2/3. Later recruitment, after one day, fully depends on the presence of vinculin and on its ability to interact with the Arp2/3. Therefore, vinculin appears to retain Arp2/3 at AJs after an initial recruitment of Arp2/3 that does not depend on vinculin-Arp2/3 interactions. Stable MCF10A cell lines expressing GFP fusions with Arp2/3 subunits that are not part of the vinculin-Arp2/3 hybrid complexes revealed a GFP signal enriched at cell-cell junctions, 1 day after cell plating (Supplementary Fig.5), thus confirming that vinculin interacts with the canonical Arp2/3 complex in MCF10A cells.

Untransformed cells stop proliferating when confluent. This phenomenon commonly called “contact inhibition of proliferation”, can be more precisely referred to as density dependence of cell cycle progression, since cells do not stop proliferating as soon as they touch each other, but rather enter less and less frequently into a new cell cycle, as the cell culture becomes denser. We observed that KO and KI cells reached a ∼30 % higher saturation density than parental MCF10A cells (Fig. 7a, Supplementary Fig.6). We then measured the number of cycling cells by estimating the % of cells incorporating EdU, an analog of thymidine incorporated in DNA during S phase. The % of cycling cells steadily decreased as a function of the cell density. KO cells behaved like parental cells, with a high cycling rate at low density and a low cycling rate at high density. However, at an intermediate density, KO cells were significantly more prone to enter into a new cell cycle than parental cells (Fig. 7b). A similar behavior was observed for KI cells (Fig. 7c), suggesting that the mere presence of vinculin was not sufficient to control contact inhibition; vinculin should also be able to interact with the Arp2/3. To confirm this point, we analyzed the density dependence of cell cycle progression MCF10A cells expressing the vinculin linker. Cells expressing the dominant-negative construct exhibited significantly increased cycling even at high cell density (Fig. 7d). In contrast, the P878A mutation in the linker abolished this increased cycling, thus reinforcing the idea that vinculin controls cell cycle progression through its ability to interact with the Arp2/3 complex. When parental cells were treated with the Arp2/3 inhibitory compound CK-666, cell cycle progression was inhibited in a dose-dependent manner (Fig. 7e). KO and KI cells, as well as MCF10A cells expressing the dominant-negative vinculin linker also displayed a dose-dependent inhibition of cell cycle progression, but required more CK-666 to achieve the same level of inhibition (Fig. 7e,f). Cell cycle progression thus appears to be inhibited by vinculin through its effect on Arp2/3 activity, in a manner similar to what occurs for membrane protrusions and persistence of single cell migration.

**Figure 7.**
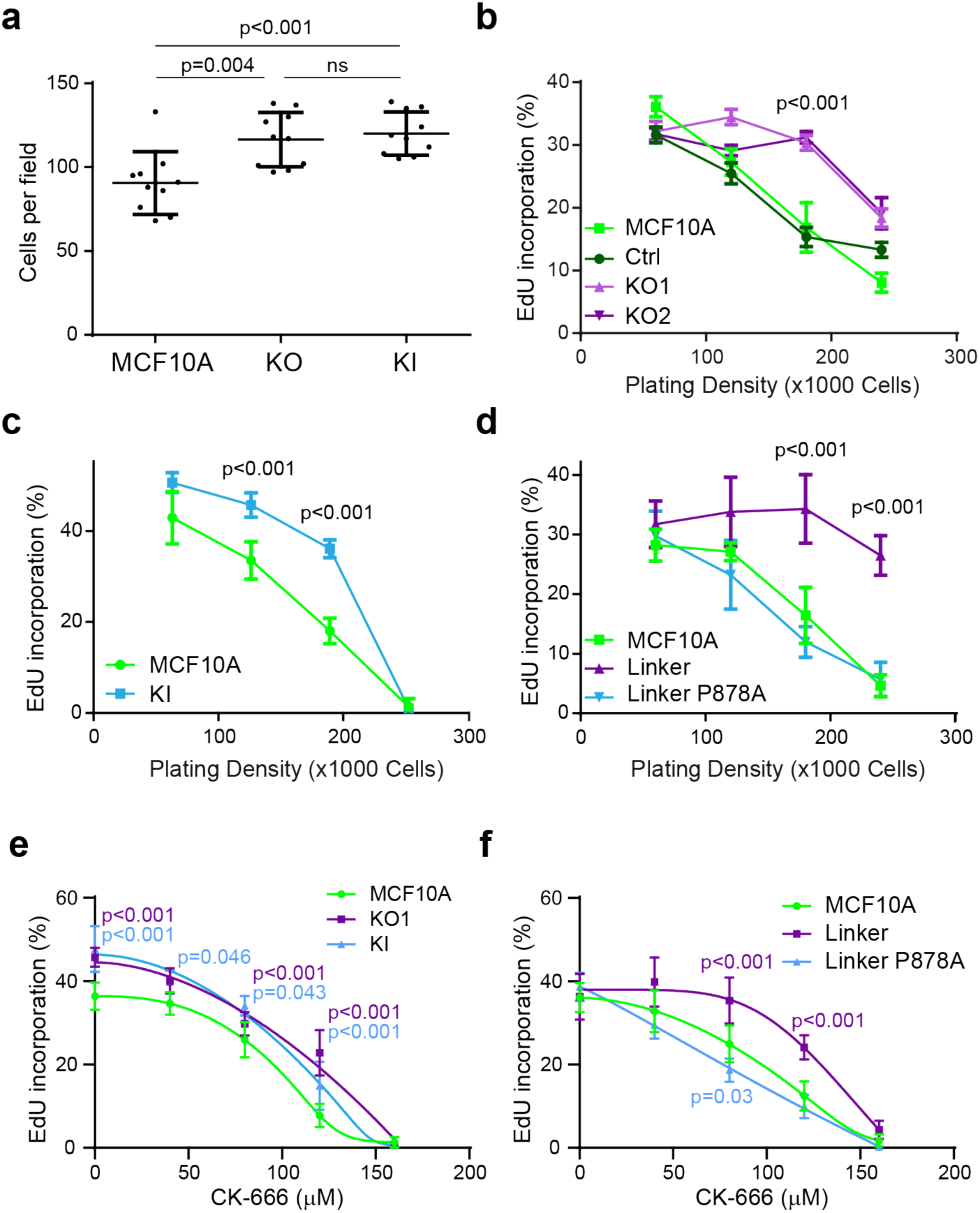
Vinculin controls cell cycle progression through its interaction with the Arp2/3 complex. **a** Saturation density of KO and KI cells. Mean ± SD, n=10, t-test. N=3, 1 representative experiment shown. **b-d** Cell cycle progression of KO (**b**), KI (**c**) and linker expressing MCF10A cells (**d**). Percentage of cells incorporating EdU is represented as a function of cell density. **e,f** Cell cycle progression of KO and KI cells (**e**), or linker expressing MCF10A cells (**f**) plated at a density of 5.10^4^ cells and treated with increasing doses of the Arp2/3 inhibitory compound CK-666. Mean ± SD, n=8 fields of views with more than 5000 cells in total, t-test. N=3, 1 representative experiment shown. p-values are shown only when both KOs or KI cells are different from both controls in **b,c,e** and when the linker is significantly different from parental MCF10A in **d,f**

## DISCUSSION

The vinculin-Arp2/3 interaction was first assumed to be a transient regulated interaction that involved the canonical Arp2/3 complex (DeMali *et al*, 2002). However, alternative assemblies of so-called vinculin-Arp2/3 hybrid complexes have then been discovered (Chorev *et al*, 2014). Since Arp2/3 subunits analyzed in vinculin immunoprecipitates by DeMali belonged to both canonical and hybrid complexes, it was not known whether the two modes of binding existed or if the vinculin-Arp2/3 interaction only involved assembly of hybrid complexes. In the present study, we unambiguously detected subunits that belong to the canonical Arp2/3 complex, but not to hybrid complexes, in vinculin immunoprecipitates or at cell-cell junctions at a time when vinculin is solely responsible for Arp2/3 recruitment. Therefore, even if these experiments do not rule out the presence of vinculin-Arp2/3 hybrid complexes in MCF10A cells, they show that the interaction of vinculin with the canonical Arp2/3 complex does exist, as suggested by the original reference. We also favor the interpretation that the roles of the vinculin-Arp2/3 interaction we report here are due to an interaction of vinculin with the canonical Arp2/3 complexes, because these functions in membrane protrusion, migration persistence and cell cycle progression were all previously ascribed to the canonical Arp2/3 complex (Wu *et al*, 2012; Suraneni *et al*, 2012; Molinie *et al*, 2019).

Our strategy to compare the phenotypes of KO and KI cells allowed us to distinguish the mechanotransducer function of vinculin, which only depends on the presence of vinculin, from the vinculin functions that require both vinculin and its interaction with Arp2/3 (Fig. 8). Among the Arp2/3-dependent functions of vinculin in MCF10A cells, we found an inhibition of branched actin assembly, membrane protrusions and cell spreading. It was reported in the original article mapping the Arp2/3 binding site on vinculin that vinculin KO mouse embryonic fibroblasts (MEFs) had impaired lamellipodia and cell spreading (DeMali *et al*, 2002). These two phenotypes were rescued by the expression of WT vinculin, but not by the P878A derivative. Our two studies thus implicate the same functions, but with opposite cellular effects.

**Figure 8.**
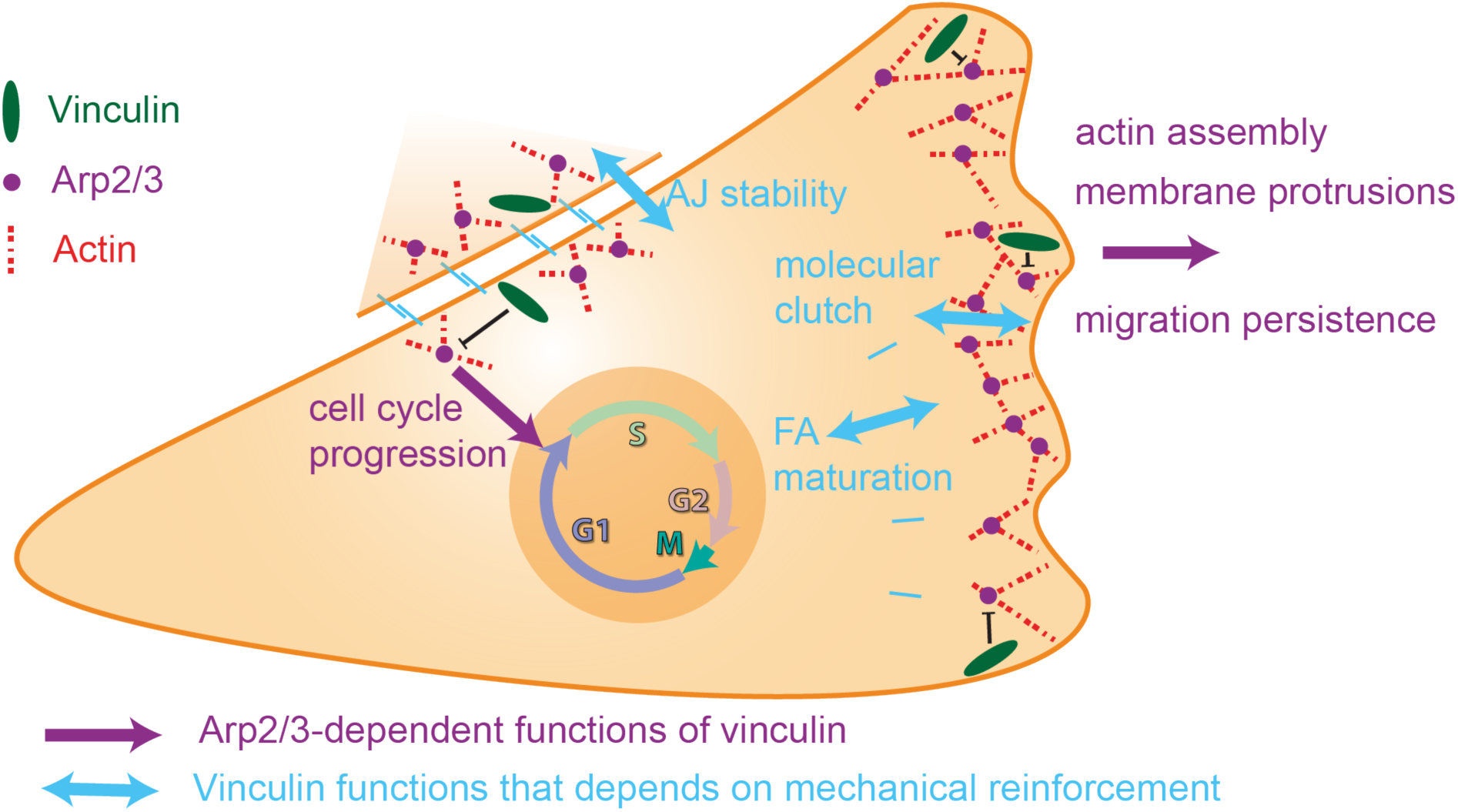
Vinculin controls cell migration and cell cycle progression through its ability to interact with the Arp2/3 complex. The vinculin-Arp2/3 interaction antagonizes branched actin polymerization in membrane protrusion and inhibits migration persistence of single cells. The actin reinforcement that vinculin provides stabilizes cell-cell junctions. Subsequent vinculin-dependent recruitment of Arp2/3 at cell-cell junctions contributes to collective migration and signals density-dependent inhibition of cell cycle progression.

Similarly, the increased migration persistence we observed in vinculin KO MCF10A cells is in contrast to the decreased persistence observed upon vinculin depletion in MEF KO cells or siRNA-treated mammary carcinoma MDA-MB-231 cells, using different assays (Thievessen *et al*, 2015; Rahman *et al*, 2016; Lee *et al*, 2019). Even if the migration phenotypes of vinculin depleted cells strongly depend on cell types and precise assay conditions, in particular 2D vs. 3D conditions (Fernández *et al*, 1993; Mierke *et al*, 2010; Thievessen *et al*, 2015), it is clear that untransformed epithelial cells MCF10A cells behave in a significantly different manner than the previously examined cell types. Another surprising observation came from the examination of the molecular clutch that connects actin polymerization at the leading edge to cell adhesions to the substratum. In fibroblasts, vinculin has been implicated in mediating the molecular clutch that connects the retrograde flow of actin to cell adhesions to the substratum (Thievessen *et al*, 2013; Hirata *et al*, 2014). Since the lamellipodia mainly consist of branched actin, one can expect that vinculin-Arp2/3 interactions could be part of the molecular clutch. In epithelial MCF10A cells, we confirm vinculin function in the molecular clutch, but exclude a role for vinculin-Arp2/3 interactions. In our vinculin KO and KI MCF10A cells, the release of branched actin inhibition can account for the increased persistence of single cell migration. Moreover, the increased efficiency of membrane protrusions in vinculin KI cells compared to KO and parental MCF10A cells fully agrees with vinculin KI cells being slightly more persistent and more spread than vinculin KO cells.

Our epithelial cell system allowed us to examine the role of vinculin at cell-cell adhesions. We found that vinculin KO MCF10A cells had dramatically decreased stability of AJs when cells were embedded into soft 3D collagen gels, highlighting the interplay between adhesions to the substratum and to the neighboring cells. Vinculin belongs to both cell adhesions, the specific incorporation into cell-cell adhesions being determined by Abl-mediated phosphorylation of Y822 (Bays *et al*, 2014). Decreased cell-cell adhesion upon vinculin KO was recently observed in mouse 4T1 breast cancer line (DeWane *et al*, 2022) and is in line with the aberrant AJs between cardiac myocytes reported in heart-specific KO of vinculin in mice (Zemljic-Harpf *et al*, 2007). Vinculin is an essential component of cell-cell junctions that allows myosin-dependent tensile forces to develop mature junctions (Twiss *et al*, 2012). KI cells do not exhibit unstable AJs and even exhibit more stable AJs than parental cells. Disabling Arp2/3 binding thus appear to increase the function of vinculin in reinforcing AJs. Increased branched actin at early cell-cell junctions can account for the increased junction stability of KI cells, if we assume that branched actin is remodeled in linear arrays for myosin-mediated contractility. Indeed, branched actin is a poor substrate for myosin motors (Muresan *et al*, 2022), but GMF and coronin proteins were shown to debranch actin networks of lamellipodia (Cai *et al*, 2008; Haynes *et al*, 2015) and might play a similar role at cell-cell junctions.

In 2D wound healing assay of MCF10A cells, vinculin KO cells were more efficient at closing the wound than parental cells, as previously reported using 4T1 cells (DeWane *et al*, 2022). KO MCF10A cells exhibited fast and directional migration towards the wound and transmitted the signal further back in the monolayer, indicating that the mechanotransduction of E-Cadherin dependent cell adhesions that vinculin provides (le Duc *et al*, 2010) is not essential to this transmission, and even rather inhibitory. These phenotypes are less prominent when vinculin is present, but impaired in its Arp2/3 interaction, indicating that the vinculin-Arp2/3 interaction is less critical for collective migration than for single cell migration.

Cadherins at AJs were found to be associated with branched actin, which pushes membranes from neighboring cells against one another to initiate cell-cell adhesion or repair unzipped membranes due to excessive tension (Efimova & Svitkina, 2018; Li *et al*, 2020, 2021; Senju *et al*, 2023). We found that vinculin was essential to recruit the Arp2/3 complex at AJs, but not at an early time point, 6 h after cell plating, where not only Arp2/3 recruitment did not depend on vinculin, but also Arp2/3 recruitment increased in KI cells. These results show that vinculin plays an essential role in Arp2/3 recruitment at AJs, but that there are also other ways to recruit it. The α-catenin molecule, which recruits vinculin at AJs, also binds to the Arp2/3 and inhibits it, but this involves a free form of α-catenin that is not bound to E-cadherin and β-catenin (Drees *et al*, 2005; Benjamin *et al*, 2010). The nucleation promoting factors, WAVE and N-WASP, recruit Arp2/3 and induce polymerization of branched actin at AJs (Kovacs *et al*, 2002; Verma *et al*, 2012; Rajput *et al*, 2013). Cortactin, which is now recognized as an Arp2/3 stabilizer of the branched junction of actin filaments (Gautreau *et al*, 2022), is also critical for Arp2/3 localization at AJs (Helwani *et al*, 2004; Han *et al*, 2014). These proteins are obvious candidates for the early vinculin-independent recruitment of Arp2/3 at AJs we observed here. However, their implication is difficult to test, since they also have a critical role in the formation and maintenance of AJs, the very structure where one would like to assess Arp2/3 recruitment.

Vinculin was found to control saturating cell density of cultures and regulate cell cycle progression as a function of cell density. This increased proliferation was observed in both KO and KI cells, indicating that this vinculin function strictly depends on its ability to interact with the Arp2/3. We previously established that cell cycle progression in untransformed cells depends on cortical branched actin (Molinie *et al*, 2019). In fact, the Arp2/3-dependent functions of vinculin uncovered here, membrane protrusions, persistence of single cell migration and cell cycle progression, were previously shown to depend on the RAC1-WAVE-Arp2/3 pathway (Dang *et al*, 2013; Molinie *et al*, 2019; Simanov *et al*, 2021). Increased protrusions, increased persistence and increased cycling observed in KO and KI MCF10A cells are phenotypes, which are all associated with increased branched actin, showing that the vinculin-Arp2/3 interaction should antagonize the formation of branched actin. Nevertheless, we found that the vinculin linker that binds the Arp2/3 in the cell was not affecting Arp2/3 activity in vitro, in the pyrene-actin assay, suggesting that additional factors or post-translational modifications are probably involved in the cell. Future work should be aimed at deciphering the precise molecular mechanisms by which vinculin antagonizes the nucleation of branched actin networks by the Arp2/3 complex.

## METHODS

### Cell Culture and Drugs

MCF10A cells were grown in DMEM/F12 medium supplemented with 5% horse serum, 20 ng/mL epidermal growth factor, 10µg/mL insulin, 100 ng/mL cholera toxin, 500 ng/mL hydrocortisone and 100U/mL penicillin. Medium and supplements were from Life Technologies and Sigma. Cells were incubated at 37°C in 5% CO2. Cells were trypsinised (12605010, Gibco) and sub-cultured every 3 days. CK-666 (182515, Sigma) was used for Arp2/3 inhibition as stated.

### Plasmids, Transfection and Isolation of Stable Cell Lines

GFP tagged proteins were expressed from a home-made vector, MXS AAVS1L SA2A Puro bGHpA EF1Flag GFP Blue SV40pA AAVS1R that was previously described (Molinie *et al*, 2019). Vinculin linker WT, P878A, ARPC1A, ARPC1B, ARPC5, ARPC5 were inserted into this plasmid in place of the Blue cassette using Fse1 and Asc1 restriction sites. The P878A mutation was generated from the WT plasmid using the Quikchange Lightning mutagenesis kit (Agilent) and primers (CCTAGGCCTCCACCAGCAGAGGAAAAGGATG, GTAGGAAAAGGAGACGACCACCTCCGGATCC).

Transfections of MCF10A cells were performed using Lipofectamine 3000 (Invitrogen). To obtain stable cell lines, MXS AAVS1 vectors were co-transfected with two TALEN constructs (Addgene #59025 and 59026) inducing a double strand break at the AAVS1 locus (González *et al*, 2014). Cells were selected with 1 µg/mL puromycin (ant-pr-1, Invivogen) and pooled if the expression was homogeneous or cloned otherwise.

### Genome Editing

To generate vinculin KO lines, MCF10A cells were transfected with a sgRNA (ATCGTGCGCGTATGAAACAC) targeting nucleotides 7 to 26 of the *VCL* coding sequence, corresponding to amino acids 3 to 9 of the vinculin protein, along with the purified Cas9 protein using the Lipofectamine CrisprMax kit (#CMAX00001, ThermoFisher). Cells were then diluted and seeded in 96-well plate at 1 cell/well. Wells containing two or more clones were not analyzed. 130 clones were screened by dot blot using anti-vinculin antibodies at 1:1000 dilution. Cells with minimal signal on dot blots were further screened using Western blot, immunostaining and sequencing to derive characterized KO clones.

To characterize the *VCL* mutations on the vinculin gene, base pairs 14-494 were amplified by PCR using DreamTaq (EP0702, ThermoFisher) for 32 cycles with annealing temperature 58°C and the (TCTGTCTCTTCGCCGGTTC, AGCCTTTTTCATGACTGCTCC) primers. The PCR product was sequenced. When several sequences overlapped, PCR products were cloned into a blunt vector using the Zero Blunt PCR cloning kit (#K270040, ThermoFisher) to sequence the two alleles independently.

To obtain the vinculin KI line, MCF10A cells were cotransfected by electroporation with a Cas9 expressing plasmid (CMV hSpCas9 bGH pA), a plasmid expressing the *ATP1A1* sgRNA (Agudelo *et al*, 2017), a pRG2(-GG) plasmid expressing the *VCL* sgRNA (GCCTCCACCACCAGAGGAAA) and a single-stranded 87 bp repair oligonucleotide (base pairs 110878-110964 in the *VCL* gene). Colonies resistant to ouabain (0.5 µM, Sigma 03125) were cloned using dilution and screened by PCR using DreamTaq (EP0702, ThermoFisher) for 32 cycles with annealing temperature 52°C and the (GGTGACGATCGAAAAAC, TATTGGCAACACAGGAACC) primers, followed by PvuII restriction.

### Antibodies

Antibodies used for immunostaining and western blots were anti-vinculin (#V9131, Sigma), anti-α-catenin (#C2081, Sigma), anti-E-cadherin (#MABT26, Merck), anti-Paxillin (#GTX125891, GeneTex), anti-ARPC2 (#07-227-I, Millipore), anti-ARPC1B (#HPA004832, Sigma), anti-ARPC3 (#HPA006550, Sigma). For immunostaining, secondary antibodies for anti-mouse-647 (#A21236, Life technologies) and anti-rabbit-405(#A34556, Life technologies) were used along with Acti-Stain 555 (Cytoskeleton).

### Immunoprecipitation and Western Blot

Two 15-cm dishes of MCF10A cells were lysed 1 day after seeding and scraped off in (50mM HEPES pH7.7, 10mM EDTA, 50mM KCl, 1mM MgCl_2_ and 1%NP-40) supplemented with a protease inhibitor cocktail (Roche, 1:10000). Cell lysates were centrifuged at 10,000xg for 10 min. The supernatant was incubated at 4°C for 1 h with GFP-Trap Agarose beads (Chromotek) and washed 4 times. Lysates and immunoprecipitates were analyzed by Western blot.

### Immunofluorescence and Image Analysis

To image cells, 15,000 cells (to obtain individual cells) or 1.5×10^6^ cells (to obtain a monolayer) were plated on 22x22 mm coverslips that were coated with fibronectin (10 µg/mL, F1141, Sigma) and fixed after 1 day (unless otherwise stated) with 4% PFA for 15 min. Cells were permeabilized in 0.2% Triton X-100 and blocked in 10% FBS in PBS. Cells were stained in a 1:200 dilution of first primary and then secondary antibodies along with Acti-stain 555. Coverslips were mounted in Dako mounting medium and imaged using an SP8 laser-scanning confocal microscope (Leica). Images were analyzed using FIJI. To measure cell spreading, cell area was quantified on images by thresholding (Triangle algorithm) in Fiji.

To quantify vinculin recruitment at AJs and FAs, the thresholded α-catenin or paxillin staining, respectively was used to generate a mask in which fluorescence intensity of vinculin was quantified. Lengths of focal adhesions were manually measured using paxillin staining. To quantify Arp2/3 and vinculin enrichment at AJs, a line was manually drawn along the junction labeled with anti-E-cadherin. Width of the line was then increased iteratively to measure total fluorescence intensity at increasing distances from the junction. Intensity at a distance n from the junction corresponded to (total intensity of line width_n_) – (total intensity of line width_n-1_). Values were finally normalized to the average fluorescence intensity at a line ≈ 12 µm from the junction, corresponding to intensity in the cytoplasm. Enrichment of E-cadherin at AJs was measured similarly using phalloidin staining as a reference for cell-cell contacts.

### Live Cell Imaging

For 2D cell migration assays, 15,000 cells were seeded 1 day prior to imaging on microslides (#80826, Ibidi) coated with fibronectin. For 3D migration of cells embedded in a collagen matrix, 15,000 cells were plated into microchambers (#80826, Ibidi) on a 3.5 mg/ml Collagen Type1 (#354236, Corning) matrix in DMEM:F12 supplemented with FBS (10%). After cells attached, another layer of collagen was added on top and cells were incubated in culture medium for 1 day before imaging. Cells were imaged on an AxioObserver Z1 microscope (Zeiss) equipped with an Orca-R2 CCD camera (Hamamatsu) and controlled by the AxioVision software (Zeiss). Images were acquired at 5 min (2D migration) or 10 min intervals (migration in 3D collagen matrix) for 24h. Cell-cell junction disassembly events were counted manually at each time point. Analysis of migration persistence was performed as previously described (Gorelik & Gautreau, 2014).

For TIRF-SIM imaging, 100,000 mCherry-actin transfected cells were plated 1 day prior to imaging on glass-bottom dishes (P35G-0.170-14-C, MatTek Corp) coated with fibronectin. To image actin flows in lamellipodia, images were acquired at 2 s intervals for 2 min using 3 phase shifted angles, each with 3 fringe patterns, on the Deltavision OMX SR (GE Healthcare) microscope. High resolution images were reconstructed and 2-color images were aligned using softWoRx (AppliedPrecision). Kymographs were generated in FIJI using manually drawn lines that followed the direction of actin retrograde flow and the Multi Kymograph tool. Protrusion speed and rearward flow are given by the tan of the angle made with the time axis in kymographs. Protrusion speeds and rearward flow values have opposite signs, but absolute values are given for clarity. The actin assembly rate is the sum of protrusion speed and rearward flow.

### Analysis of wound healing experiments

80,000 cells were plated on microslides (#80826, Ibidi) coated with fibronectin within inserts (#80209, Ibidi) one day before the experiment. Inserts were removed, and timelapse acquisitions were performed at 10 min intervals during 24h on an AxioObserver Z1 microscope (Zeiss) equipped with a 10x objective, an Orca-Flash4 v3 camera (Hamamatsu) and controlled by the MicroManager 2.0 software. Fields focused on only one edge of the wound were acquired, and average leading-edge positions perpendicular to the axis of the wound gave leading edge progression. The velocity field was obtained by PIV (Petitjean *et al*, 2010) using the PIVlab software package (Thielicke & Sonntag, 2021) for Matlab (MathWorks). The window size was set to 32 pixels, i.e. 23.75 µm with a 0.75 overlap between windows. A time sliding window averaging velocity fields over 40 minutes (equal to 4 frames) was used. Spurious vectors were filtered-out by their amplitude and were replaced by interpolated velocities from neighboring vectors. The local order parameter was calculated using a previously published code (Deforet *et al*, 2012).

### Cell Cycle and Proliferation

To quantify saturation density, 2×10^6^ cells were plated on fibronectin coated cover slips in 6-well plates. Cells were fixed 4 days after seeding and nuclei were stained with DAPI. To perform the EdU incorporation assay, cells were seeded on fibronectin-coated coverslips (12 mm) for 1 day. Cells were incubated with 10 µM EdU for 1 h prior to fixation in 4% PFA for 15 min and permeabilized in 1% Triton X-100 for 5 min. EdU was labeled with the Alexa Fluor 488 Click-iT EdU Imaging Kit (#C10337, Thermo Fisher Scientific) according to the manufacturer instructions and nuclei were labeled with DAPI (Thermo Fisher Scientific). Images were acquired on an inverted microscope (Olympus IX83) using a 20x objective (NA 0.5) equipped with an Orca-Flash4.0 V3 camera (Hamamatsu), controlled by Micro-Manager 2.0 and analyzed using a custom script in FIJI to count DAPI- or EdU-positive cells. The percentage of cells in S-phase was scored as the ratio of EdU-positive nuclei/DAPI-stained nuclei in segmented images. For each condition, at least 5000 cells were counted.

### Statistical Analysis

For all t-tests, populations were first tested for Gaussian distribution using a Shapiro-Wilk Test with a α-value of 0.05. If both populations were Gaussian, difference between means was tested using Welch’s T-test and if one or both of the populations were non-Gaussian, the difference between means was tested using a Mann–Whitney U test. focal adhesion size, protrusion speed and frequency of cell junction disassembly were tested with a one-tailed distribution as the lower limit of measurement was close to a lower bound of 0. All other t-tests assumed a two-tailed distribution. Analysis of migration persistence was performed as previously described (Gorelik & Gautreau, 2014). Exponential decay and plateau fit 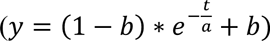 were performed for all individual cells. Coefficients were then compared using one-way ANOVA. Statistical analysis was performed in R using linear mixed-effect models to take into account the resampling of the same statistical unit as previously described (Polesskaya *et al*, 2022).

## Supporting information

Supplementary informations

## Acknowledgments

This work was supported by grants from Agence Nationale de la Recherche (ANR-20-CE13-0016, ANR-22-CE44-0006 and ANR-22-CE13-0041), Fondation ARC pour la Recherche sur le Cancer (ARC PJA 2021 060003815), and from Institut National du Cancer (INCA_16712).

## Author contributions

J.J. performed most of the experiments and analyses, and participated in writing of the manuscript. A.I.F. performed wound healing experiments and some EdU assays. P.S. supervised analysis of wound healing experiments. D.Y.G. generated the vinculin KI cell line. C.L.C. supervised actin polymerization assays performed by H.W. A.P. and S.N.R. performed the co-IP experiment. S.R. and A.M.G. have jointly supervised the work and wrote the manuscript together.

## REFERENCES

Abella JVG, Galloni C, Pernier J, Barry DJ, Kjær S, Carlier M-F & Way M (2016) Isoform diversity in the Arp2/3 complex determines actin filament dynamics. Nature Cell Biology 18: 76–86

Agudelo D, Duringer A, Bozoyan L, Huard CC, Carter S, Loehr J, Synodinou D, Drouin M, Salsman J, Dellaire G, et al (2017) Marker-free coselection for CRISPR-driven genome editing in human cells. Nat Methods 14: 615–620

Alexandrova AY, Arnold K, Schaub S, Vasiliev JM, Meister J-J, Bershadsky AD & Verkhovsky AB (2008) Comparative dynamics of retrograde actin flow and focal adhesions: formation of nascent adhesions triggers transition from fast to slow flow. PLoS ONE 3: e3234

Atherton P, Stutchbury B, Jethwa D & Ballestrem C (2016) Mechanosensitive components of integrin adhesions: Role of vinculin. Exp Cell Res 343: 21–27

Auernheimer V, Lautscham LA, Leidenberger M, Friedrich O, Kappes B, Fabry B & Goldmann WH (2015) Vinculin phosphorylation at residues Y100 and Y1065 is required for cellular force transmission. J Cell Sci 128: 3435–3443

Bays JL & DeMali KA (2017) Vinculin in cell–cell and cell–matrix adhesions. Cellular and Molecular Life Sciences 74: 2999–3009

Bays JL, Peng X, Tolbert CE, Guilluy C, Angell AE, Pan Y, Superfine R, Burridge K & DeMali KA (2014) Vinculin phosphorylation differentially regulates mechanotransduction at cell–cell and cell–matrix adhesions. The Journal of Cell Biology 205: 251–263

Benjamin JM, Kwiatkowski AV, Yang C, Korobova F, Pokutta S, Svitkina T, Weis WI & Nelson WJ (2010) αE-catenin regulates actin dynamics independently of cadherin-mediated cell–cell adhesion. J Cell Biol 189: 339–352

Bieling P & Rottner K (2023) From WRC to Arp2/3: Collective molecular mechanisms of branched actin network assembly. Curr Opin Cell Biol 80: 102156

Blanchoin L, Boujemaa-Paterski R, Sykes C & Plastino J (2014) Actin Dynamics, Architecture, and Mechanics in Cell Motility. Physiological Reviews 94: 235–263

Borgon RA, Vonrhein C, Bricogne G, Bois PRJ & Izard T (2004) Crystal Structure of Human Vinculin. Structure 12: 1189–1197

Boujemaa-Paterski R, Martins B, Eibauer M, Beals CT, Geiger B & Medalia O (2020) Talin-activated vinculin interacts with branched actin networks to initiate bundles. Elife 9: e53990

Cai L, Makhov AM, Schafer DA & Bear JE (2008) Coronin 1B Antagonizes Cortactin and Remodels Arp2/3-Containing Actin Branches in Lamellipodia. Cell 134: 828–842

Chorev DS, Moscovitz O, Geiger B & Sharon M (2014) Regulation of focal adhesion formation by a vinculin-Arp2/3 hybrid complex. Nature communications 5: 1–11

Ciobanasu C, Faivre B & Le Clainche C (2014) Actomyosin-dependent formation of the mechanosensitive talin-vinculin complex reinforces actin anchoring. Nature communications 5: 3095

Dang I, Gorelik R, Sousa-Blin C, Derivery E, Guérin C, Linkner J, Nemethova M, Dumortier JG, Giger FA, Chipysheva TA, et al (2013) Inhibitory signalling to the Arp2/3 complex steers cell migration. Nature 503: 281–284

Deforet M, Parrini MC, Petitjean L, Biondini M, Buguin A, Camonis J & Silberzan P (2012) Automated velocity mapping of migrating cell populations (AVeMap). Nature Methods 9: 1081–1083

del Rio A, Perez-Jimenez R, Liu R, Roca-Cusachs P, Fernandez JM & Sheetz MP (2009) Stretching single talin rod molecules activates vinculin binding. Science 323: 638–641

DeMali KA, Barlow CA & Burridge K (2002) Recruitment of the Arp2/3 complex to vinculin: coupling membrane protrusion to matrix adhesion. The Journal of Cell Biology 159: 881–891

DePasquale JA & Izzard CS (1991) Accumulation of talin in nodes at the edge of the lamellipodium and separate incorporation into adhesion plaques at focal contacts in fibroblasts. The J Cell Biology 113: 1351–1359

DeWane G, Cronin NM, Dawson LW, Heidema C & DeMali KA (2022) Vinculin Y822 is an important determinant of ligand binding. J Cell Sci

Drees F, Pokutta S, Yamada S, Nelson WJ & Weis WI (2005) Alpha-catenin is a molecular switch that binds E-cadherin-beta-catenin and regulates actin-filament assembly. Cell 123: 903–915

Efimova N & Svitkina TM (2018) Branched actin networks push against each other at adherens junctions to maintain cell–cell adhesion. J Cell Biol 217: 1827–1845

Fernández JR, Geiger B, Salomon D & Ben-Ze’ev A (1993) Suppression of vinculin expression by antisense transfection confers changes in cell morphology, motility, and anchorage-dependent growth of 3T3 cells. The J Cell Biology 122: 1285–1294

Garrido-Casado M, Asensio-Juárez G & Vicente-Manzanares M (2021) Nonmuscle Myosin II Regulation Directs Its Multiple Roles in Cell Migration and Division. Annu Rev Cell Dev Biology 37: 1–26

Gautreau AM, Fregoso FE, Simanov G & Dominguez R (2022) Nucleation, stabilization, and disassembly of branched actin networks. Trends Cell Biol 32: 421–432

González F, Zhu Z, Shi Z-D, Lelli K, Verma N, Li QV & Huangfu D (2014) An iCRISPR platform for rapid, multiplexable, and inducible genome editing in human pluripotent stem cells. Cell stem cell 15: 215–226

Gorelik R & Gautreau A (2014) Quantitative and unbiased analysis of directional persistence in cell migration. Nature Protocols 9: 1931–1943

Han SP, Gambin Y, Gomez GA, Verma S, Giles N, Michael M, Wu SK, Guo Z, Johnston W, Sierecki E, et al (2014) Cortactin Scaffolds Arp2/3 and WAVE2 at the Epithelial Zonula Adherens. The Journal of biological chemistry 289: 7764–7775

Haynes EM, Asokan SB, King SJ, Johnson HE, Haugh JM & Bear JE (2015) GMF controls branched actin content and lamellipodial retraction in fibroblasts. The Journal of Cell Biology 209: 803–812

Helwani FM, Kovacs EM, Paterson AD, Verma S, Ali RG, Fanning AS, Weed SA & Yap AS (2004) Cortactin is necessary for E-cadherin–mediated contact formation and actin reorganization. J Cell Biology 164: 899–910

Hirata H, Tatsumi H, Lim CT & Sokabe M (2014) Force-dependent vinculin binding to talin in live cells: a crucial step in anchoring the actin cytoskeleton to focal adhesions. Am J Physiol-Cell Physiol 306: C607–C620

Humphries JD, Wang P, Streuli C, Geiger B, Humphries MJ & Ballestrem C (2007) Vinculin controls focal adhesion formation by direct interactions with talin and actin. The J Cell Biology 179: 1043–1057

Jasin M & Haber JE (2016) The democratization of gene editing: Insights from site-specific cleavage and double-strand break repair. Dna Repair 44: 6–16

Kadota M, Yang HH, Gomez B, Sato M, Clifford RJ, Meerzaman D, Dunn BK, Wakefield LM & Lee MP (2010) Delineating Genetic Alterations for Tumor Progression in the MCF10A Series of Breast Cancer Cell Lines. Plos One 5: e9201

Kovacs EM, Goodwin M, Ali RG, Paterson AD & Yap AS (2002) Cadherin-directed actin assembly: E-cadherin physically associates with the Arp2/3 complex to direct actin assembly in nascent adhesive contacts. Current Biology 12: 379–382

Krause M & Gautreau A (2014) Steering cell migration: lamellipodium dynamics and the regulation of directional persistence. Nature Reviews Mol Cell Biol 15: 577–590

Le Clainche C & Carlier M-F (2008) Regulation of actin assembly associated with protrusion and adhesion in cell migration. Physiological Reviews 88: 489–513

Le Clainche C, Dwivedi SP, Didry D & Carlier M-F (2010) Vinculin Is a Dually Regulated Actin Filament Barbed End-capping and Side-binding Protein. The Journal of biological chemistry 285: 23420–23432

le Duc Q, Shi Q, Blonk I, Sonnenberg A, Wang N, Leckband D & Rooij J de (2010) Vinculin potentiates E-cadherin mechanosensing and is recruited to actin-anchored sites within adherens junctions in a myosin II–dependent manner. The Journal of Cell Biology 189: 1107–1115

Lee HT, Sharek L, O’Brien ET, Urbina FL, Gupton SL, Superfine R, Burridge K & Campbell SL (2019) Vinculin and metavinculin exhibit distinct effects on focal adhesion properties, cell migration, and mechanotransduction. PLoS ONE 14: e0221962

Leerberg JM, Gomez GA, Verma S, Moussa EJ, Wu SK, Priya R, Hoffman BD, Grashoff C, Schwartz MA & Yap AS (2014) Tension-Sensitive Actin Assembly Supports Contractility at the Epithelial Zonula Adherens. Curr Biology 24: 1689–1699

Li JXH, Tang VW, Boateng KA & Brieher WM (2021) Cadherin puncta are interdigitated dynamic actin protrusions necessary for stable cadherin adhesion. Proc National Acad Sci 118: e2023510118

Li JXH, Tang VW & Brieher WM (2020) Actin protrusions push at apical junctions to maintain E-cadherin adhesion. Proceedings of the National Academy of Sciences of the United States of America 117: 432–438

Mierke CT, Kollmannsberger P, Zitterbart DP, Diez G, Koch TM, Marg S, Ziegler WH, Goldmann WH & Fabry B (2010) Vinculin Facilitates Cell Invasion into Three-dimensional Collagen Matrices*. J Biological Chem 285: 13121–13130

Moese S, Selbach M, Brinkmann V, Karlas A, Haimovich B, Backert S & Meyer TF (2007) The Helicobacter pylori CagA protein disrupts matrix adhesion of gastric epithelial cells by dephosphorylation of vinculin. Cell Microbiol 9: 1148–1161

Molinie N, Rubtsova SN, Fokin A, Visweshwaran SP, Rocques N, Polesskaya A, Schnitzler A, Vacher S, Denisov EV, Tashireva LA, et al (2019) Cortical branched actin determines cell cycle progression. Cell Research 29: 432–445

Muresan CG, Sun ZG, Yadav V, Tabatabai AP, Lanier L, Kim JH, Kim T & Murrell MP (2022) F-actin architecture determines constraints on myosin thick filament motion. Nat Commun 13: 7008

Papalazarou V & Machesky LM (2021) The cell pushes back: The Arp2/3 complex is a key orchestrator of cellular responses to environmental forces. Curr Opin Cell Biol 68: 37–44

Petitjean L, Reffay M, Grasland-Mongrain E, Poujade M, Ladoux B, Buguin A & Silberzan P (2010) Velocity Fields in a Collectively Migrating Epithelium. Biophys J 98: 1790–1800

Pizarro-Cerdá J, Chorev DS, Geiger B & Cossart P (2016) The Diverse Family of Arp2/3 Complexes. Trends in Cell Biology: 1–8

Polesskaya A, Boutillon A, Yang S, Wang Y, Romero S, Liu Y, Lavielle M, Molinie N, Rocques N, Fokin A, et al (2022) Restrained activation of CYFIP2-containing WAVE complexes controls membrane protrusions and cell migration. Biorxiv: 2020.07.02.184655

Pollard TD (2016) Actin and Actin-Binding Proteins. Cold Spring Harbor perspectives in biology 8: a018226–18

Pollard TD & Cooper JA (2009) Actin, a central player in cell shape and movement. Science 326: 1208–1212

Poujade M, Grasland-Mongrain E, Hertzog A, Jouanneau J, Chavrier P, Ladoux B, Buguin A & Silberzan P (2007) Collective migration of an epithelial monolayer in response to a model wound. Proceedings of the National Academy of Sciences 104: 15988–15993

Rahman A, Carey SP, Kraning-Rush CM, Goldblatt ZE, Bordeleau F, Lampi MC, Lin DY, García AJ & Reinhart-King CA (2016) Vinculin regulates directionality and cell polarity in two- and three-dimensional matrix and three-dimensional microtrack migration. Mol Biology Cell 27: 1431–1441

Rajput C, Kini V, Smith M, Yazbeck P, Chavez A, Schmidt T, Zhang W, Knezevic N, Komarova Y & Mehta D (2013) Neural Wiskott-Aldrich Syndrome Protein (N-WASP)-mediated p120-Catenin Interaction with Arp2-Actin Complex Stabilizes Endothelial Adherens Junctions* ♦. J Biological Chem 288: 4241–4250

Seddiki R, Narayana GHNS, Strale P-O, Balcioglu HE, Peyret G, Yao M, Le AP, Lim CT, Yan J, Ladoux B, et al (2018) Force-dependent binding of vinculin to α-catenin regulates cell-cell contact stability and collective cell behavior. Molecular Biology of the Cell 29: 380–388

Senju Y, Mushtaq T, Vihinen H, Manninen A, Saarikangas J, Ven K, Engel U, Varjosalo M, Jokitalo E & Lappalainen P (2023) Actin-rich lamellipodia-like protrusions contribute to the integrity of epithelial cell–cell junctions. J Biol Chem 299: 104571

Simanov G, Dang I, Fokin AI, Oguievetskaia K, Campanacci V, Cherfils J & Gautreau AM (2021) Arpin Regulates Migration Persistence by Interacting with Both Tankyrases and the Arp2/3 Complex. Int J Mol Sci 22: 4115

Soule HD, Maloney TM, Wolman SR, Peterson WD, Brenz R, McGrath CM, Russo J, Pauley RJ, Jones RF & Brooks SC (1990) Isolation and characterization of a spontaneously immortalized human breast epithelial cell line, MCF-10. Cancer Research 50: 6075–6086

Subauste MC, Pertz O, Adamson ED, Turner CE, Junger S & Hahn KM (2004) Vinculin modulation of paxillin–FAK interactions regulates ERK to control survival and motility. The J Cell Biology 165: 371–381

Suraneni P, Rubinstein B, Unruh JR, Durnin M, Hanein D & Li R (2012) The Arp2/3 complex is required for lamellipodia extension and directional fibroblast cell migration. The Journal of Cell Biology 197: 239–251

Thielicke W & Sonntag R (2021) Particle Image Velocimetry for MATLAB: Accuracy and enhanced algorithms in PIVlab. J Open Res Softw 9: 12

Thievessen I, Fakhri N, Steinwachs J, Kraus V, McIsaac RS, Gao L, Chen B, Baird MA, Davidson MW, Betzig E, et al (2015) Vinculin is required for cell polarization, migration, and extracellular matrix remodeling in 3D collagen. The FASEB J 29: 4555–4567

Thievessen I, Thompson PM, Berlemont S, Plevock KM, Plotnikov SV, Zemljic-Harpf A, Ross RS, Davidson MW, Danuser G, Campbell SL, et al (2013) Vinculin–actin interaction couples actin retrograde flow to focal adhesions, but is dispensable for focal adhesion growth. J Cell Biol 202: 163–177

Twiss F, le Duc Q, Horst SVD, Tabdili H, Krogt GVD, Wang N, Rehmann H, Huveneers S, Leckband DE & Rooij JD (2012) Vinculin-dependent Cadherin mechanosensing regulates efficient epithelial barrier formation. Biology open 1: 1128–1140

Verma S, Han SP, Michael M, Gomez GA, Yang Z, Teasdale RD, Ratheesh A, Kovacs EM, Ali RG & Yap AS (2012) A WAVE2-Arp2/3 actin nucleator apparatus supports junctional tension at the epithelial zonula adherens. Molecular Biology of the Cell 23: 4601–4610

Vigouroux C, Henriot V & Le Clainche C (2020) Talin dissociates from RIAM and associates to vinculin sequentially in response to the actomyosin force. Nat Commun 11: 3116

Worsham MJ, Pals G, Schouten JP, Miller F, Tiwari N, Spaendonk R van & Wolman SR (2005) High-resolution mapping of molecular events associated with immortalization, transformation, and progression to breast cancer in the MCF10 model. Breast Cancer Research and Treatment 96: 177–186

Wu C, Asokan SB, Berginski ME, Haynes EM, Sharpless NE, Griffith JD, Gomez SM & Bear JE (2012) Arp2/3 is critical for lamellipodia and response to extracellular matrix cues but is dispensable for chemotaxis. Cell 148: 973–987

Yao M, Goult BT, Chen H, Cong P, Sheetz MP & Yan J (2014a) Mechanical activation of vinculin binding to talin locks talin in an unfolded conformation. Nature Reviews Mol Cell Biol 4: 259–7

Yao M, Qiu W, Liu R, Efremov AK, Cong P, Seddiki R, Payre M, Lim CT, Ladoux B, Mège R-M, et al (2014b) Force-dependent conformational switch of α-catenin controls vinculin binding. Nature communications 5: 4525

Zemljic-Harpf AE, Miller JC, Henderson SA, Wright AT, Manso AM, Elsherif L, Dalton ND, Thor AK, Perkins GA, McCulloch AD, et al (2007) Cardiac-Myocyte-Specific Excision of the Vinculin Gene Disrupts Cellular Junctions, Causing Sudden Death or Dilated Cardiomyopathy. Mol Cell Biol 27: 7522–7537

Zhang Z, Izaguirre G, Lin S-Y, Lee HY, Schaefer E & Haimovich B (2004) The Phosphorylation of Vinculin on Tyrosine Residues 100 and 1065, Mediated by Src Kinases, Affects Cell Spreading. Mol Biology Cell 15: 4234–4247

