## Supplementary informations for "Vinculin-Arp2/3 Interaction Inhibits Branched Actin Assembly to Control Cell Migration and Cell Cycle Progression"

**SUPPLEMENTARY INFORMATION**

**Supplementary Figures ……………….....................… 2**

**Legends to Supplementary Movies ………….…...…. 8**

### SUPPLEMENTARY FIGURES

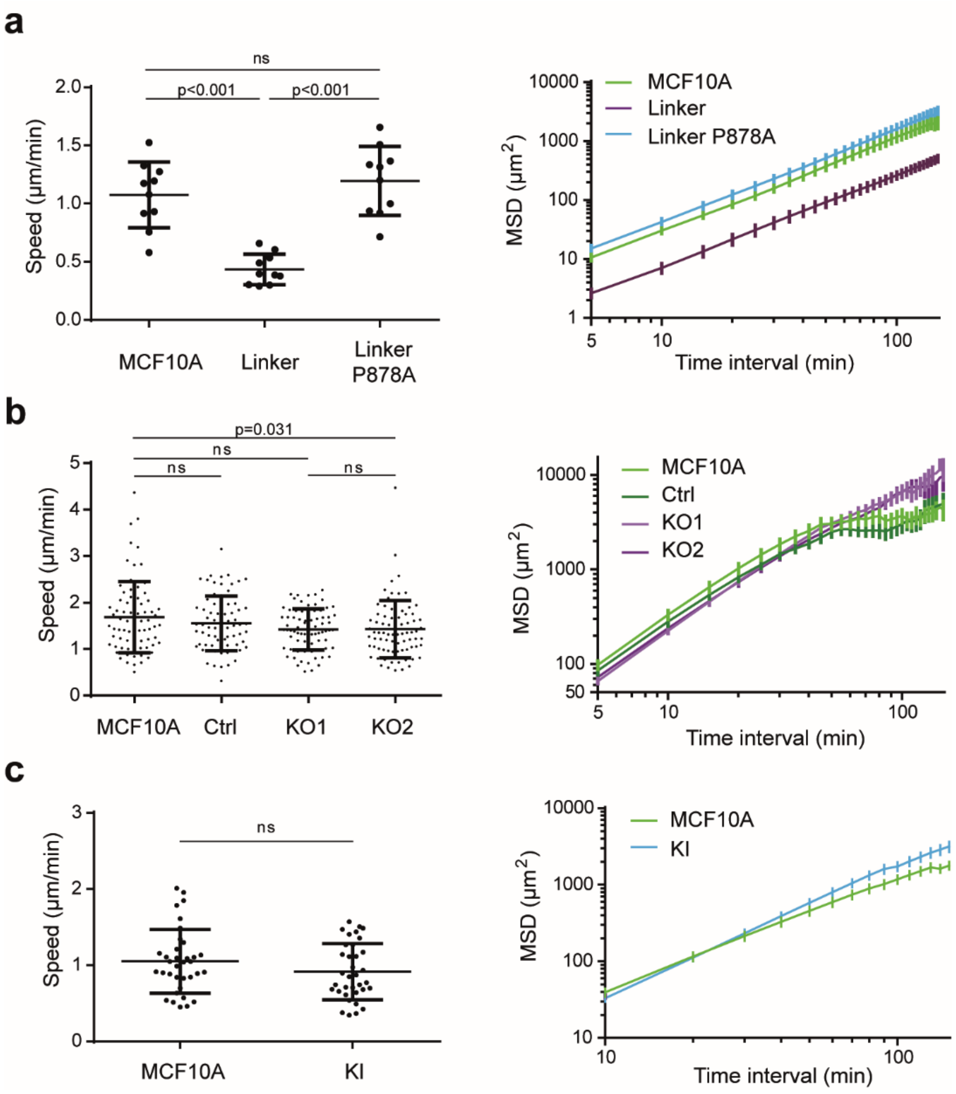


**Supplementary Figure 1. Speed and Mean Square Displacement (MSD). a** MCF10A cells expressing the vinculin linker (n=10). **b** KO cells (n=74). **c** KI cells (n=35). Mean ± SD, t-test, N=3, 1 representative experiment shown.

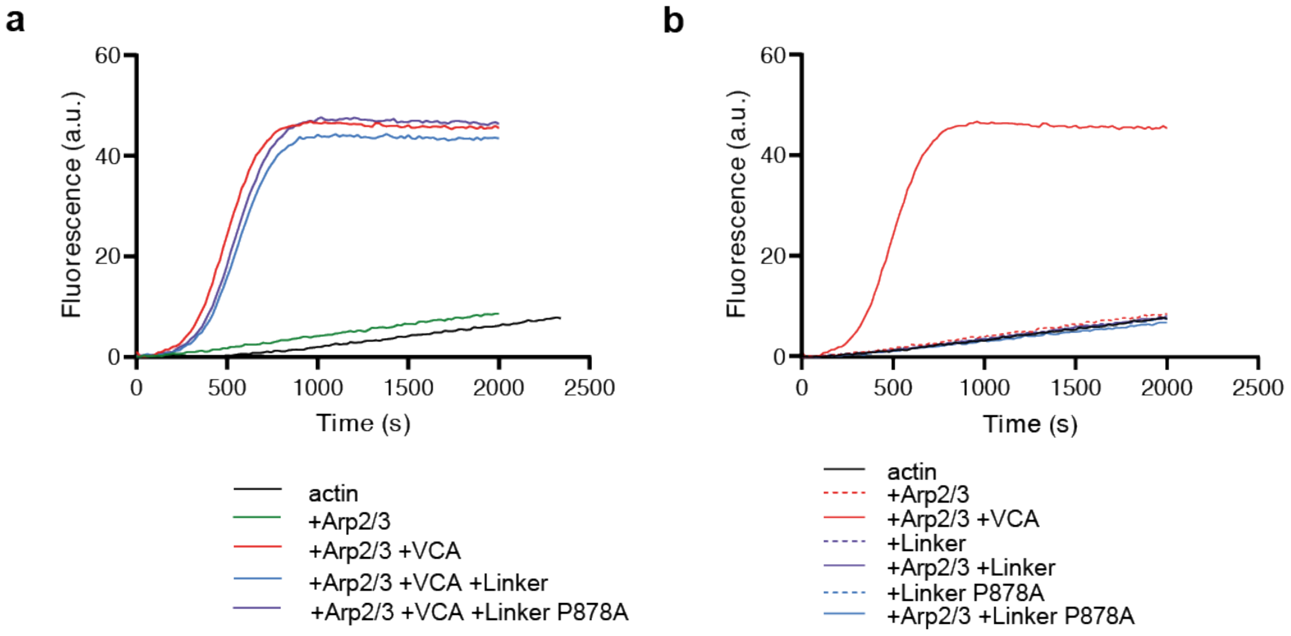


**Supplementary Figure 2. Pyrene-actin polymerization assays. a** The vinculin linker does not regulate Arp2/3 activity in vitro. **b** The vinculin linker does not affect actin polymerization in vitro. Conditions: 1.5 µM of 10% pyrenyl-labelled actin, 20 nM Arp2/3, 250 nM VCA and 1 µM vinculin linker or P878A linker as indicated. All these curves were acquired the same day and the Arp2/3 + VCA is replotted in the two panels for comparison.

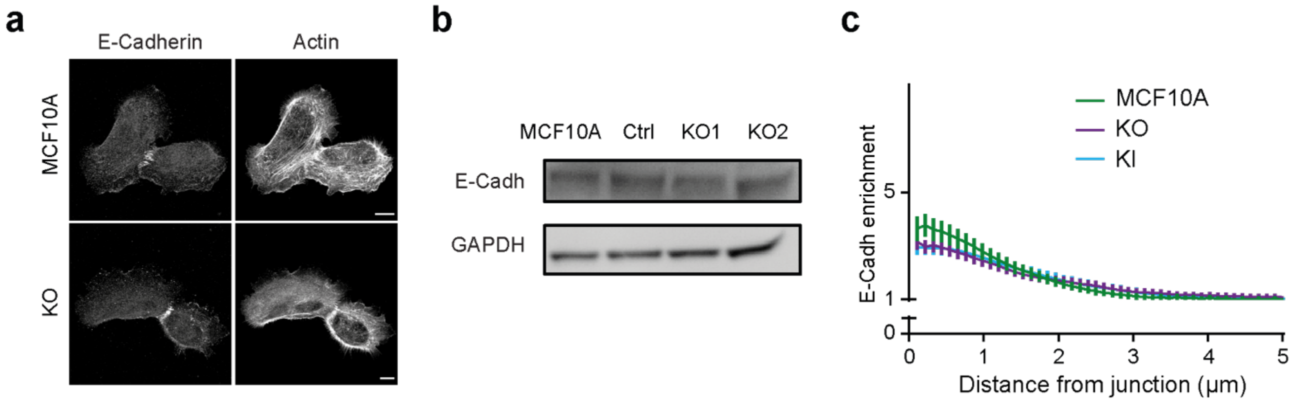


**Supplementary Figure 3. E-Cadherin in KO and KI clones. (a)** Staining of E-cadherin and F-actin using phalloidin of parental and KO cells, 1 day after plating on glass coverslips. Scale bars 10 µm. **(b)** Western blot analysis of KO clones. **(c)** Enrichment of E-Cadherin at cell-cell junctions of parental, KO and KI cell monolayers, 1 day after plating. Mean ± SD at each distance are plotted. N=3, 1 representative experiment shown.

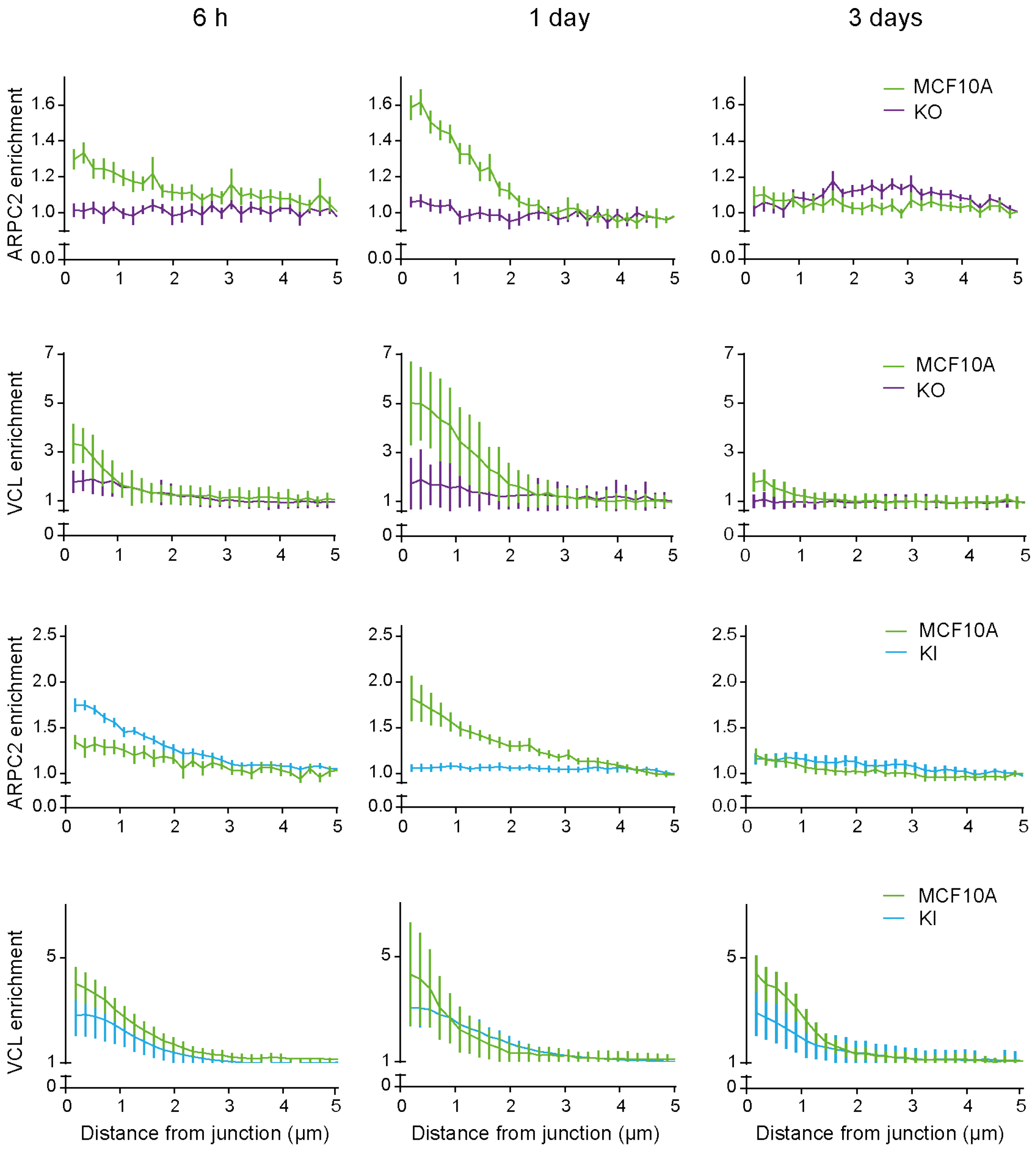


**Supplementary Figure 4. Enrichment of ARPC2 and vinculin at cell-cell junctions of KO and KI cells.** Normalized intensity of staining plotted against distance from junctions, 6 hours, 1 day and 3 days after plating. Mean ± SD at each distance are plotted. n=10, N=3, 1 representative experiment shown.

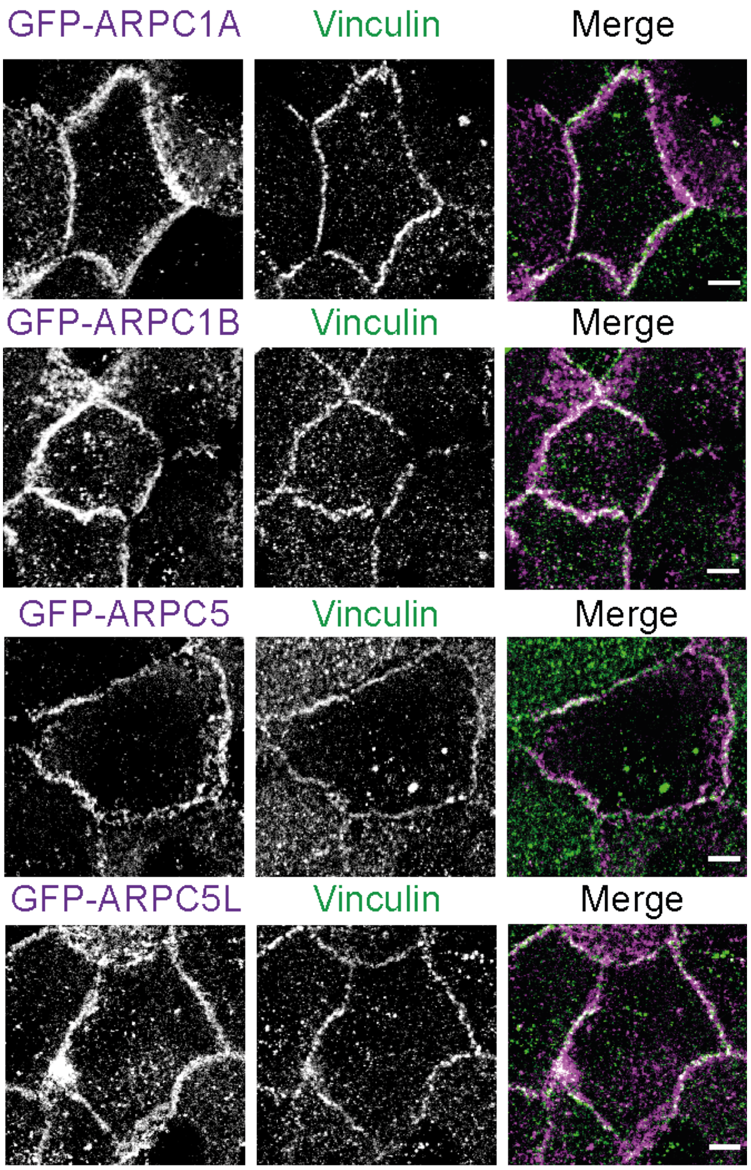


**Supplementary Figure 5. Localisation of various Arp2/3 subunits at cell-cell junctions.** Stable MCF10A clones expressing the indicated GFP fusion proteins were fixed 1 day after plating and stained with vinculin antibodies. Max z-projection from confocal microscopy. Scale bars 5 µm.

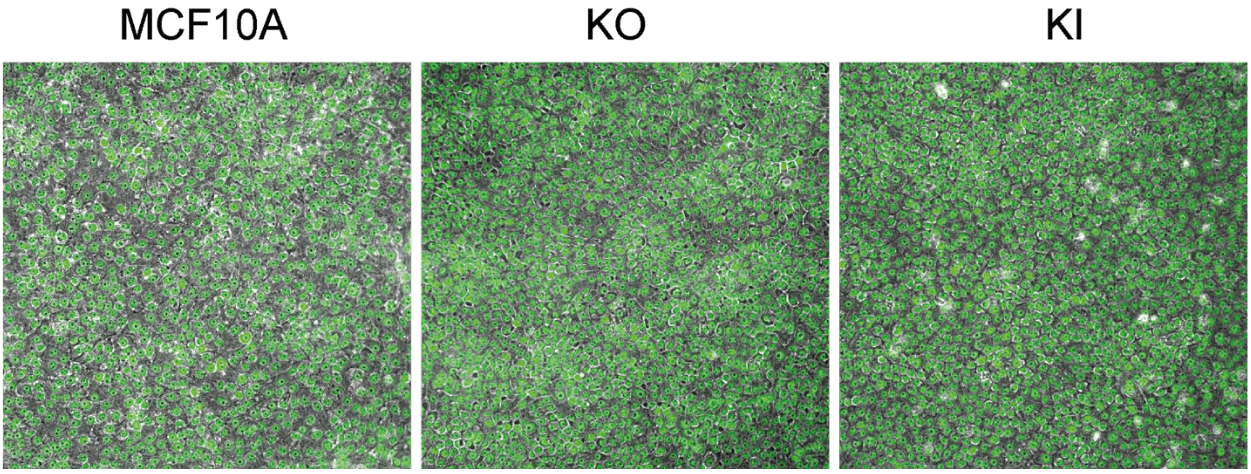


**Supplementary Figure 6. Saturation density of KO and KI cells.** DAPI staining (green) was overlaid onto phase contrast images.

### LEGENDS TO SUPPLEMENTARY MOVIES

**Supplementary Movie 1, related to figure 1d. Single cell migration of MCF10A cells expressing the vinculin linker.** Cell tracks are superimposed on phase contrast images. Scale bar: 10 µm.

**Supplementary Movie 2, related to figure 1h. Actin dynamics at membrane protrusions of MCF10A cells expressing the vinculin linker.** Stable MCF10A cells expressing the WT or the P878A linker were transiently transfected with a plasmid encoding mCherry-actin. TIRF-SIM. Scale bar: 1 µm.

**Supplementary Movie 3, related to figure 3. Single cell migration of KO and KI cells.** Cell tracks are superimposed on phase contrast images. Scale bar: 10 µm.

**Supplementary Movie 4, related to figure 3. Actin dynamics at membrane protrusions of KO and KI cells.** MCF10A, KO2 and KI cells were transiently transfected with a plasmid encoding mCherry-actin. TIRF-SIM. Scale bars, 2 µm.

**Supplementary Movie 5, related to figure 4e.** MCF10A and KO2 cells were sandwiched between two collagen gels and imaged with phase contrast optics. Scale bar: 15 µm.

**Supplementary Movie 6, related to figure 4f.** MCF10A and KI cells were sandwiched between two collagen gels and imaged with phase contrast optics. Scale bar: 15 µm.

**Supplementary Movie 7, related to figure 5. Collective migration of KO and KI cells upon wound healing.** Phase contrast time lapse of MCF10A, KO2 and KI cells acquired immediately after lifting an insert. Scale 100 µm.
